# Neural representations of aversive value encoding in pain catastrophizers

**DOI:** 10.1101/279992

**Authors:** Christopher A. Brown, Abeer F. Almarzouki, Richard J. Brown, Anthony K. P. Jones

## Abstract

Chronic pain is exacerbated by maladaptive cognition such as pain catastrophizing (PC). Biomarkers of PC mechanisms may aid precision medicine for chronic pain. Here, we investigate EEG biomarkers using mass univariate and multivariate (machine learning) approaches. We test theoretical notions that PC results from a combination of augmented aversive-value encoding (“magnification”) and persistent expectations of pain (“rumination”). Healthy individuals with high or low levels of PC underwent an experimental pain model involving nociceptive laser stimuli preceded by cues predicting forthcoming pain intensity. Analysis of EEG acquired during the cue and laser stimulation provided event-related potentials (ERPs) identifying spatially and temporally-extended neural representations associated with pain catastrophizing. Specifically, differential neural responses to cues predicting high vs. low intensity pain (i.e. aversive value encoding) were larger in the high PC group, largely originating from mid-cingulate and superior parietal cortex. Multivariate spatiotemporal EEG patterns evoked from cues with high aversive value selectively and significantly differentiated the high PC from low PC group (64.6% classification accuracy). Regression analyses revealed that neural patterns classifying groups could be partially predicted (R^2^ = 28%) from those neural patterns classifying the aversive value of cues. In contrast, behavioural and EEG analyses did not provide evidence that PC modifies more persistent effects of prior expectation on pain perception and nociceptive responses. These findings support the hypothesis of magnification of aversive value encoding but not persistent expression of expectation in pain catastrophizers. Multivariate patterns of aversive value encoding provide promising biomarkers of maladaptive cognitive responses to chronic pain that have future potential for psychological treatment development and clinical stratification.

## 1 Introduction

Across a range of physical health conditions, cognitive factors are known to impact on outcomes (Edwards et al., 2011; Ottaviani et al., 2016; Trick et al., 2016). While there has been substantial interest in identifying biomarkers that increase the precision of psychiatric classification and predict outcomes in mental health disorders (Singh and Rose, 2009), more research is needed into biomarkers of cognitive risk factors for physical health conditions. For example, chronic pain is highly prevalent in the population, estimated to be from 19% to 50% depending on survey methods and definitions of severity (Croft et al., 2010). However, treatments for chronic pain are poorly targeted to underlying mechanisms (Jones and Brown, 2017), partly due to a lack of viable biomarkers. Suggested biomarkers have ranged from salivary cortisol (van Aken et al., 2018) to BOLD signals (Woo and Wager, 2015), but research has not specifically focussed on markers of cognitive risk factors.

A maladaptive cognitive trait that is widely studied in relation to chronic pain is pain catastrophizing (PC). PC has been defined as “an exaggerated negative mental set brought to bear during actual or anticipated pain experience” (Sullivan et al., 1995). PC predicts the severity of chronic pain (Edwards et al., 2011) and physical dysfunction above and beyond the effects of concurrent depression (Arnow et al., 2011). While the importance of PC in chronic pain is rarely disputed, its sub-component mechanisms have not so far been clearly defined.

Sub-components of PC can be described in both psychological and neurobiological terms. The most common characterisation of PC has been in terms of the three subscales of the Pain Catastrophizing Scale (PCS (Sullivan et al., 1995)) which recognises augmentation of the aversive value of pain (“magnification”), perseverative thinking about pain (“rumination”) and deficits in coping ability (“helplessness”). Although historically there are alternative conceptualisations of PC (reviewed in (Neblett, 2017)), in the present work we utilise the tripartite concept underlying the PCS. However, neurobiological research has tended to focus on the neural correlates of PC as a unitary construct. EEG and fMRI studies in chronic pain patients have found greater activation of the secondary somatosensory cortex to non-painful stimuli (Vase et al., 2012) and painful stimuli (Gracely et al., 2004) in pain catastrophizers, as well as greater activation of anterior cingulate cortex in both healthy volunteers (Seminowicz and Davis, 2006) and fibromyalgia patients (Gracely et al., 2004). Further research provided more precise mechanistic insights by identifying anticipatory deficits in lateral prefrontal cortex activity in chronic pain patients with greater levels of PC (Brown et al., 2014; Loggia et al., 2015), suggestive of a failure of the top-down inhibitory control provided by this brain region (Lorenz et al., 2003), which can be remedied through psychological intervention (Brown and Jones, 2013). This provides a basis for further investigation of clinically viable biomarkers of PC.

One approach is to investigate PC in pain-free individuals, in order to avoid the potential confound of chronic pain symptoms. In this study, we acquired scalp EEG in pain-free individuals with high and low levels of PC in order to characterise PC in terms of two hypothesised neural processes: aversive value encoding (“magnification”) and perseverative expectation effects (“rumination”). These processes were operationalised with respect to the *transient* (magnification) and *persistent* (rumination) effects of expectancy cues on the temporal dynamics of neural responses as participants anticipated and experienced experimental (laser) pain stimuli. This approach to perseverative expectation builds on our previous research showing that expectancy effects on nociception and pain persist even if initial expectancy cues are followed by contrary information indicating no threat of pain (the so-called ‘prior expectancy effect’ (Almarzouki et al., 2017)). Here we tested the hypothesis of a more persistent expectancy effect (i.e. augmented nociceptive processing and reported pain) in pain catastrophizers. Secondly we tested an alternative hypothesis in which magnified cue processing occurs in pain catastrophizers more transiently without affecting subsequent pain processing.

Our analyses utilised standard mass univariate as well as multivariate analysis of event-related potentials (ERPs) in order to identify spatiotemporal patterns that classify high from low PC groups, as a step towards biomarker development. Specifically, we used Multivariate Pattern Analysis (MVPA), which applies machine-learning algorithms to neuroimaging data. In recent years, MVPA has provided predictive measures of pain at the single individual level (Brodersen et al., 2012; Wager et al., 2013) and has been applied to ERPs to classify patients with psychiatric disorders (Taylor et al., 2017); here, we provide the first attempt to apply this methodology to classifying PC.

## 2 Methods

### 2.1 Study design

A single session 2×2x2 mixed design was used, with group (high, low catastrophizing) as the between-participants variable and two within-participants variables each with two levels: Expectancy (“prior high” and “low”) and laser heat intensity (“medium” and “low”); in addition to these conditions, additional trials were randomised into the procedure that served to reinforce pain expectancy responses to high cues by following these cues with high intensity laser stimuli (figure 1). The study consisted of a psychophysics test, a training procedure, and then the experiment proper, which took place over the course of two blocks of stimulation.

**Figure 1.**
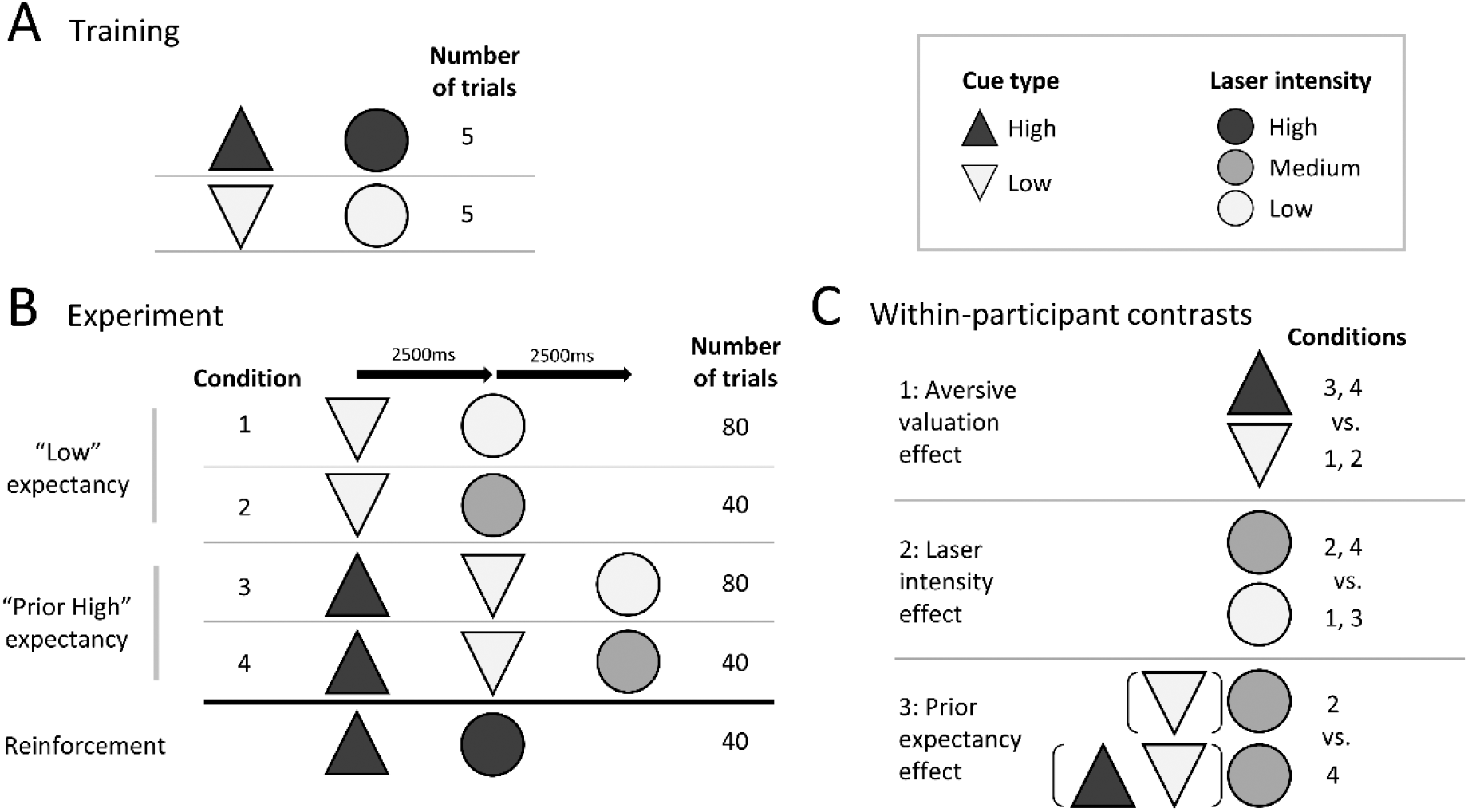
Conditioning procedure, experimental conditions and contrasts for analysis. A) A brief training procedure involved randomised trials of high and low intensity laser stimuli (pain and non-painful respectively), each of which was preceded by a visual cue that reliably predicted the laser intensity. B) The experiment consisted of randomised trials of conditions 1 to 4 plus reinforcement trials. The conditions differed according to the presentation of visual cues consisting of upward and downward-pointing arrows (shown here as triangles), and variably intense laser stimuli (represented as circles). In conditions 1 and 2, “low” cues were presented, which most of the time (two-thirds, condition 2) led to a low intensity laser stimulus, while the other one-third of trials (condition 3) led to a medium intensity laser stimulus that they had previously not been trained to expect (and were not informed might occur). Conditions 3 and 4 consisted of a “high” cue that was replaced after 2.5s by a “low” cue. Following this after another 2.5s, two-thirds of trials (condition 3) led to a low intensity laser stimulus, while the other one-third of trials (condition 4) led to a medium intensity laser stimulus, in a similar way to the contingency in conditions 1 and 2 (which are identical except for the lack of a prior “high” cue). Condition 5 acted as a reinforcement of their prior expectation from training, i.e. that upward arrows lead to high intensity pain. On every trial, after the laser stimulus participants were asked rate the pain intensity on a 0 – 10 numerical rating scale. C) For EEG analysis, three contrasts were derived from comparisons of conditions 1-4. Contrast 1 was the aversive valuation contrast on cue processing, namely EEG responses to the “high” cues (conditions 3 and 4) vs. the “low” cues (conditions 1 and 2). Contrast 2 was the laser intensity contrast of medium intensity stimuli (conditions 2 and 4) vs. low intensity stimuli (conditions 1 and 3). Contrast 3 was the prior pain expectancy contrast on laser stimulus processing, namely medium intensity laser stimulus processing from condition 4 vs. condition 2.

### 2.2 Power calculation

A large effect size (Cohen’s d > 0.8) was anticipated based on the recruitment strategy: study power was maximised by recruiting participants scoring in the upper and lower quartile of the Pain Catastrophizing Scale (PCS – see below), providing a large group separation on this measure (for illustration purposes, the *measured* (not *expected*) Cohen’s d effect size was 1.92 – see supplementary table 1) that was expected to translate to large effect sizes in the outcomes of interest (pain ratings and EEG responses). The recruitment target of 52 participants was exceeded but due to exclusions (see Supplementary Methods) the final sample size was 34, providing power to detect effect sizes on group differences on the order of Cohen’s d = 1.0 with 80% power and an alpha of 0.05.

### 2.3 Ethics and recruitment

Ethical approval was obtained from North West Nine Research Ethics Committee in the United Kingdom. Volunteers were mainly recruited through The University of Manchester. Participants received an honorarium of £10 per hour, in addition to travel expenses.

### 2.4 Participants: Screening

All participants described themselves as above 18, right handed, free from pain, neurological illness, morbid psychiatric illness, peripheral vascular disease, ischemic heart disease, chronic skin disease (e.g. eczema, psoriasis) and hypertension not controlled by medication.

Volunteers were screened by scores on the self-report Pain Catastrophizing Scale (PCS) (Sullivan et al., 1995), which they completed prior to recruitment by email or through an online survey tool. The PCS is a 13-item questionnaire relating to thoughts and feelings about pain, with 5-point Likert-scale response categories ranging from zero (“not at all”) to five (“all the time”). The questionnaire consists of separate subscales for rumination, magnification, and helplessness, which can be calculated separately or combined to form a total score.

### 2.5 Participants: Grouping

Participants with total PCS scores in the upper (hitherto referred to as the “high PC” group) and lower quartile (“low PC”) of the PCS were identified according to the score ranges in the scale manual (Sullivan et al., 1995). These participants were invited to participate in the study. Participants were asked to repeat the PCS upon attending the laboratory to ensure the answers given during screening were reliable.

### 2.6 Study procedures

#### 2.6.1 Participant expectations prior to the experiment

While the PCS provided a general (trait) measure of expected distress from painful situations, it did not specify participants’ expected distress from the particular situation of this experiment. To check whether participants with high PCS scores were indeed anticipating the laser stimuli in this experiment to be more distressing that the low PCS group, following Brown et al. (Brown et al., 2008b), participants were asked to rate their expected distress using nine items from the profile of mood states (POMS) scale (*sad, angry, discouraged, hopeless, hostile, irritable, tense, anxious* and *worried*). Specifically, they were asked to rate the extent to which they expected to experience each of these emotions while experiencing the laser heat pulses during the experiment, rating each on a 5-point Likert scale from 0 (‘‘not at all”) to 4 (‘‘very much”). The sum of these items was taken as a measure of anticipated emotional distress as done previously (Brown et al., 2008b; Sullivan et al., 2001).

#### 2.6.2 Extraneous participant variables

In addition to questionnaire measurement of PC, participants completed (prior to the experiment) a number of other questionnaires to characterise them psychologically; these data were used to shed light on the specificity of the groups to the PC construct. These additional instruments were: Depression (Patient Health Questionnaire 9, PHQ-9) (Spitzer et al., 2001), somatic symptoms (Patient Health Questionnaire 15, PHQ-15) (Kroenke et al., 2002), State and Trait Anxiety (State Trait Anxiety Inventory, STAI) (Spielberger et al., 2010) which includes separate subscales for state and trait anxiety, and the Fear of Pain Questionnaire (FPQ) with subscales for minor pain, severe pain and medical pain (Roelofs et al., 2005).

#### 2.6.3 Laser stimulation and psychophysics

All participants received “high” (moderately painful), “medium” (pain threshold) and “low” (non-painful) intensity laser stimuli during the experiment. Laser stimuli were used due to their high selectivity to A-delta and C nociceptive fibers (Meyer et al., 1976), and were administered using a thulium laser with a beam diameter of 6 mm and pulse duration of 100 milliseconds to the dorsal surface of the right forearm. The pulses were systematically moved around the skin surface to avoid skin sensitisation, damage, and habituation. Participants wore protective safety goggles throughout the study.

We sought to provoke roughly equivalent levels of painful and non-painful stimuli across individuals. Hence, laser intensities were individually calibrated using a psychophysics test prior to the experiment proper. In this test, participants rated the intensity of each stimulus on a 0 to 10 numerical pain rating scale (NRS) in which level 4/10 was defined as pain threshold. Three additional points on the scale were defined as anchors to enable consistency across participants: level 3/10 (low-intensity stimulus) was described to the volunteer as hot, but not painful; level 5/10 (medium-intensity stimulus) was described as a low and ignorable painful sensation; level 7/10 (high-intensity stimulus) was described as moderately painful and not easy to ignore. To find these levels for each participant, the intensity of the stimuli was gradually increased starting from an imperceptible level and progressing to the moderately painful level (7/10), as decided by the participant. This was repeated three times. At the end of the test, the three levels were selected based on averaged scores from the three runs.

#### 2.6.4 Training procedure

Participants were trained to calibrate their expectations of pain intensity in relation to two types of visual cues that would be presented prior to laser stimulation during the experiment. A brief training procedure (figure 1A) involved presentations of upward or downward arrows that predicted (after 2.5s) the occurrence of high and low intensity laser stimuli respectively. There were 10 trials in total consisting of five trials of each cue/laser pairing (randomised order). Participants were not made aware that they would also be presented with medium intensity laser stimuli during the subsequent experiment.

#### 2.6.5 Main experiment

Participants were informed that, on each trial of the main experiment, the trial would start with an initial cue (a downward or upward arrow) that would inform them about the subsequent stimulus intensity. In addition, they were informed that, unlike during training, the arrows might change before the laser stimulus was delivered and that, in this case, it was only the arrow directly preceding each laser stimulus that was accurate in predicting its intensity.

There were four experimental conditions (figure 1B), which varied according to the expectations created by different cue stimuli and the consistency between those cues and the laser stimulus that followed. Regarding expectancy cue conditions, all experimental trials presented a low intensity cue (downward arrow) followed by a laser stimulus that the participant was asked to rate. In conditions 1 and 2 (“low” expectancy conditions), only a “low” cue was presented, followed by the laser stimulus 2.5s later. In conditions 3 and 4 (“prior high” expectancy conditions), the “low” cue was preceded by a “high” cue, which was delivered 2.5s earlier; the laser stimulus also occurred 2.5s after the “low” cues on these trials. Each cue was presented for 2s. This design enabled us to compare EEG responses to the “high” cues (conditions 3 and 4) vs. the “low” cues (conditions 1 and 2) to identify the effect of aversive valuation of the cues (figure 1C).

Regarding the orthogonal second within-participant factor in the design, namely laser stimulus intensity, in conditions 1 and 3 the intensity of the laser stimulus was consistent with the “low” cue on most (two thirds) of the trials; in the remaining third of trials (conditions 2 and 4) a medium intensity laser stimulus was delivered. This enabled the second major comparison which was with regarding to post-stimulus processing between medium intensity stimuli (conditions 2 and 4) vs. low intensity stimuli (conditions 1 and 3) to identify the effects of laser intensity (N.B. the numbers of trials each intensity were matched for analysis purposes - see analysis section). The third comparison, also on post-stimulus processing, involved only medium intensity stimuli from condition 4 vs. condition 2 to identify the prior pain expectancy effect.

A fifth trial type was also used (but for the analysis, not regarded as an experimental condition of interest), in which only “high” cues were delivered followed by a high intensity laser stimulus 2.5s later. These trials were designed to ensure that the “high” cue was perceived as a meaningful predictor of a high intensity stimulus in conditions 3 and 4. These trials were not included in the EEG analyses. The five trial types were delivered in random order, across two blocks of 140 trials each, with each block containing the same proportion of stimuli from each condition. The two blocks differed according to the instructions given, with participants being asked to focus on the painfulness of the stimuli in block 1 and to identify the location of the stimulus in block 2. However, to test the hypotheses in the current analysis, this block difference was not of interest; to account for any variance in the results as a results of task or time effects over blocks, block was included as a (nuisance) factor in the statistical models.

#### 2.6.6 Behavioural measures

The primary behavioural outcome was the volunteers’ self-reported pain ratings (especially, for medium intensity stimuli) and their modulation by cues. To record this, on every trial, after the laser stimulus participants were prompted to rate the pain intensity on a 0 – 10 numerical rating scale (NRS) via appearance of the scale on the computer screen three seconds after the laser pulse. Volunteers reported their pain using a button pad.

#### 2.6.7 Post-experiment manipulation check

After completion of the experiment we checked that participants were engaging with the visual cues. Three questions were asked: (i) During the task, how much did you focus on the direction of the arrows? (ii) How accurately did the arrow cues predict the intensity of the pain that followed? (iii) When rating the intensity of the pain, how much was your rating based on the direction of the preceding arrow cue? In each case, participants were asked to select an answer ranging from “not at all” to “all the time”. Participants who reported ignoring the cues completely or most of the time were excluded from the analysis.

#### 2.6.8 EEG acquisition parameters

Electroencephalography (EEG) was acquired during the main experiment from 59 Ag/AgCl surface electrodes attached to an elastic cap placed in accordance with the extended international 10-20 system (BrainVision ActiCap combined with a Neuroscan head box and amplifier system). Band-pass filters were set at DC to 100Hz with a sampling rate of 500Hz. Electrodes were referenced to the ipsilateral (right) earlobe and later (during analysis) re-referenced to the common average. In addition to the 59 scalp channels, the horizontal and vertical electro-oculograms (EOG) were measured for detection of eye-movement and blink artefacts.

### 2.7 EEG data pre-processing

EEG data pre-processing was performed using EEGLAB version 13.1.1 (Delorme and Makeig, 2004). Continuous data were initially low pass filtered at 45Hz to exclude electrical noise. Data were then segmented into epochs including from -5500ms preceding the laser stimulus (including from -500ms pre-cue) to 2000ms post-stimulus. Data containing excessive eye movement or muscular artefact were rejected by a quasi-automated procedure: noisy channels and epochs were identified by calculating their normalised variance and then manually rejected or retained by visual confirmation. Independent component analysis (ICA) based on the Infomax ICA algorithm (Bell and Sejnowski, 1995) was run on the clean data excluding bad channels using the ‘runica’ function in EEGLAB. ICA components were visually inspected and bad components rejected. Such components were identified if matching the description of either (a) high (neuronally infeasible) amplitude signals with characteristic topographic (frontal) distributions matching that of blinks or horizontal eye movements, or (b) topographies characteristic of single electrode noise but that were not frequently occurring enough to be considered as bad channels. The median number of components rejected was 8 (range 4 to 11), commonly consisting of 3-4 eye movement components and 5-6 channel noise components. After ICA correction, bad channels previously identified by visual inspection were then replaced by spherical spline interpolation of neighbouring electrodes. Data were then re-referenced to the average of 59 channels (excluding reference and ocular channels). ERPs were calculated for each subject and condition by averaging epochs.

### 2.8 EEG statistical analysis

Here we summarise our approach to the statistical analysis of the EEG data and refer to a more detailed description of the steps taken in the supplementary materials. Our analyses focussed on identifying within-subject effects and then characterising the groups in terms of these effects. The three within-subjects effects of interest were as follows. Contrast 1: the aversive cue valuation effect (High cue (conditions 3 and 4) > Low cue (conditions 1 and 2)); Contrast 2: the main effect of laser stimulus intensity (Medium (conditions 2 and 4) > Low (conditions 1 and 3)); Contrast 3: the prior expectancy effect on post-laser stimulus nociceptive processing of medium intensity stimuli only (Prior High expectation (condition 4) > Low expectation (condition 2)). For each effect, and then in further comparing effects between groups, three EEG statistical analysis approaches were used: (1) mass univariate analysis (MUA) of the sensor data, (2) MUA of source data, and (3) multivariate pattern analysis (MVPA) of sensor data.

The univariate and multivariate analyses had different but complementary goals. MUA is sensitive to mean differences in neural activity within localised regions; hence it can be used to test for interactions in amplitude of evoked responses, in specific spatiotemporal regions, between groups and conditions from mixed designs. Hence we used MUA to analyse the localisation (in sensor and source space) of amplitude differences for the effects of interest. In contrast, MVPA ignores average univariate effects to focus on neural *patterns*. One advantage of applying MVPA to EEG (as opposed to fMRI) data is the ability to utilise high-resolution temporal information. MVPA is sensitive to distributed coding of neural information (Jimura and Poldrack, 2012) across space and time, providing spatiotemporal patterns that classify groups and conditions. We depart from commonly used methods that consider time points as independent of each other (e.g. applying searchlight analysis over time (King and Dehaene, 2014)) by assuming that brain representations are coded across spatial and temporal dimensions in an integrated fashion. This spatiotemporal approach to MVPA has been successfully applied recently to the classification of patients with schizophrenia (Taylor et al., 2017) and provides algorithmic and statistical efficiency. Furthermore, group-level MVPA was preferred here over subject-level MVPA for within-subjects contrasts, as it has been recently shown to provide more consistent and interpretable results across subjects (Gilron et al., 2017).

The details of both the MUA and MVPA are in supplementary materials. Briefly, MUA sensor analyses were based on the sensor-by-time approach using the Statistical Parametric Mapping software (SPM12, www.fil.ion.ucl.ac.uk/spm) (Litvak and Friston, 2008). General Linear Models (GLMs) were estimated at the group level to produce three F-contrasts of interest: (1) the main within-subjects effect of interest (aversive valuation, pain intensity or prior expectancy effect), (2) the main effect of Group (between-subjects factor of interest), (2) the interaction of these between and within-subjects factors. Source analysis (in SPM12) then focussed on time windows that showed statistically significant effects in the sensor analysis. Sensor and source results are reported based on cluster-level significance.

MVPA analyses (see supplementary materials) involved learning classifiers (specifically, a Gaussian process classifier (GPC) (Rasmussen and Williams, 2004)) on sensor images at the 2^nd^ (group) level using the Pattern Recognition for Neuroimaging Toolbox (PRoNTo) (Schrouff et al., 2013b). This identifies patterns differentiating levels of both within-subject factors of interest (cue type and prior expectation) and between groups. Performance of models were tested using a leave-one-subject-out-per-group cross-validation scheme, while statistical significance of classification accuracy was assessed using permutation tests with 10,000 sets of randomised target labels. Further analyses were conducted on weight maps resulting from MVPA classification using the approach of *a posteriori* weight summarization (Schrouff et al., 2013a). We also transformed the classifier weights back into activation patterns, providing an interpretable time-course, as done previously (Haufe et al., 2014). Lastly, we investigated the possibility of shared representations contributing to both group and condition classifications (i.e. the representational similarity (Kriegeskorte, 2008)) via regression of weight matrices outputted from the different classifications, with robust statistics calculated via bootstrapping.

## 3 Results

In this study we focus on addressing two key issues. Firstly, is pain catastrophizing characterised by magnified aversive value encoding? Secondly, do pain catastrophizers show evidence of more persistent pain expectancy (the ‘prior expectancy effect’ (Almarzouki et al., 2017)), which would follow if pain catastrophizing is characterised by perseverative processing of pain cues (I.e., rumination)? In addition to analysing expectancy effects on pain behaviourally, EEG analyses detail the spatiotemporal location and pattern of within-subject effects of cue valuation (visual evoked potential (VEP) responses to cues predicting high vs. low pain) and the effects of prior pain expectancy on the laser-evoked potential (LEP). We then proceed to compare these within-subject effects to group effects (high vs. low PC groups).

### 3.1 Participants

Analyses were conducted on data from 16 high pain catatrophisers (“high PC”) and 18 low pain catatrophisers (“low PC”) (see supplementary results for screening, recruitment and retention numbers). The high PC group had a mean (±SD) total Pain Catastrophizing Scale (PCS) score of 33.6 (±3.8) compared to the low PC group with a mean score of 4.6 (±3.1). There was no statistically significant age difference between the two groups and groups were exactly gender balanced: high PC group (8 female, 8 male; Age M = 26.6, SD = 10.8); low PC group (9 female, 9 male; Age M = 22.6, SD = 4.7). All participants reported not being on any regular medications. The characteristics of each participant are provided in more detail in supplementary table 1. There were also group differences in some extraneous psychological variables (see supplementary table 2); of particular relevance is a significant difference in state anxiety; high PC group: 38.6 (±8.6), low PC group 29.7 (±8.8), *p* < .002 (uncorrected), Cohen’s *d* = 1.02. Group differences were also present with medium effect sizes, but were not statistically significant, for fear of medical pain and for trait anxiety.

### 3.2 Group characteristics: Behavioural results

The behavioural analyses tested for group differences in dependent variables related to pain perception (pain threshold and pain ratings). To summarise, the high (vs. low) PC group were expecting to be more distressed by the laser pain stimuli and had a lower pain threshold, but they did not show evidence of a larger “prior pain expectancy” effect (supplementary tables 1, 4 and 5).

More precisely, the high PC group reported significantly greater expectations of emotional distress from the laser stimuli (M=9.7, SD=4.3) compared to the low PC group (M=4.4, SD=4.5; *t* = 3.48, *p* < .001, Cohen’s *d* = 1.03), consistent with the view that catastrophic cognitions alter pain-related expectations. Furthermore, the mean laser energy used to reach pain threshold was lower in the high PC group (M=29.6, SD=7.2) compared to the low PC group (M=34.1, SD=5.7) with a large effect size (Cohen’s *d* = 0.67) but in an independent samples *t*-test the difference was only marginally significant at an uncorrected *p* value of 0.051. Both groups were found to score roughly equally for all “manipulation check” variables, such that there was no evidence of a difference in the extent to which the two groups reported attending to anticipation cues, believing in the accuracy of the cues or being influenced by the cues.

These results are consistent with an analysis of pain ratings in response to laser stimuli during the experiment and the effect of anticipation cues on these ratings. Specifically, pain ratings did not differ overall between groups, which is as expected given that the laser energy used for each participant was adjusted to their individual pain perception. Regarding prior expectancy effects on pain (i.e. main effect of presence vs. absence of the first “high” expectancy cue), although there was a statistically significant within-participant cue effect on pain ratings (*F*(1,32) = 37.7, *p* < .001, *η*_*p*_^2^ = 0.54, supplementary table 5), there was no interaction between group and this cue effect (p=0.938), indicating that the prior pain expectancy effect does not differ between groups. Comparing the effect size of the prior expectancy effect in each group separately, the effects are similar: *η*_*p*_^2^ = 0.49 in the high PC group and *η*_*p*_^2^ = 0.60 in the low PC group.

Further ANOVA results show that changes in laser intensity reliably modulated pain perception (*F*(1,32) = 161.7, *p* <.001, *η*_*p*_^2^ = 0.83). There was no main effect of block on pain perception and block did not interact with expectancy, but did interact with intensity (*F*(1,32) = 8.69, *p* = .006, *η_p_^2^* = 0.21), with lower intensity ratings in block 2, although this effect was not of interest to the analysis. Group did not interact with these within-subject factors.

In addition to the hypothesis-driven analyses so far described, further exploratory cross-correlation statistics (Spearman’s rank, 2-tailed) across some relevant variables recorded are reported in supplementary table 3. These are reported across all participants (n=34, or 33 in cases of missing data), but not within groups as these would not be robust considering the small sample size. These did not reveal any relationships that were in addition to those already reflected in the t-test statistics comparing the PC groups, except for expected correlations between subscales of the same measurement, e.g. PCS subscales).

### 3.3 Is pain catastrophizing characterised by magnified aversive value encoding?

To answer this question, we first investigated neural activity related to the encoding of aversive value from the visual cues using Contrast 1 of High cue > Low cue (figure 1C), independently of group. The results (figure 2) identified specific post-cue latencies and cortical sources that maximally differed between conditions (univariate analyses), in addition to spatiotemporal ERP patterns predicting the cue type (MVPA analyses). MVPA is particularly sensitive to distributed coding across space and time and is therefore useful for identifying spatiotemporal patterns classifying conditions. We then proceeded to investigate group differences (figure 3) to address whether these differences were related to within-subject aversive valuation effects. In summary of the results, within-subject aversive valuation effects were partially distinct, but also partially overlapping in space and time with group effects; furthermore, multivariate spatiotemporal patterns classifying cue conditions were able to predict group differences to a moderate degree (figure 4).

**Figure 2:**
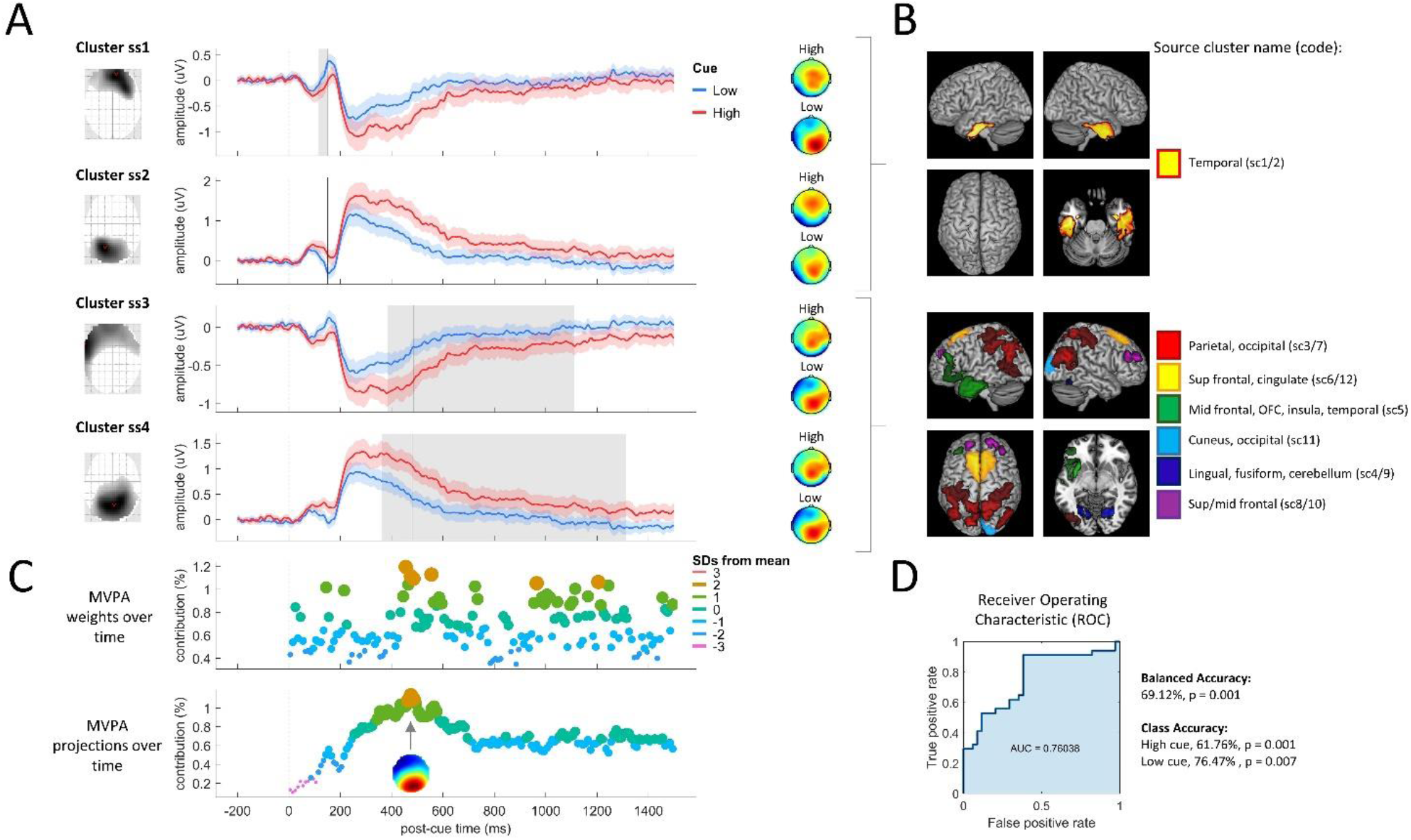
Aversive value encoding during cue processing (Contrast 1, N=34). A. Univariate sensor analysis found four statistical spatiotemporal clusters that differed in the contrast of High>Low cue. Clusters ss1 and ss2 are positive and negative polarities of the same ERP components during the early latency time window at 116ms to 152ms. Clusters ss3 and ss4 are positive and negative components during the late latency period of 362ms to 1312ms. Left: SPM glass brain statistical maps; these are the Maximum Intensity Projections (MIPs) for each cluster across all time points, thresholded to include only statistically-significant voxels; the view is the same as for EEG topographic maps, i.e. top is anterior scalp. Middle: grand average waveforms and 95% CIs; waveforms were generated by multiplying the (thresholded) SPM cluster’s MIP with the raw topographic image for each subject and condition, averaging over the remaining voxels for each time point separately; grey indicates cluster extent, black vertical line indicates latency with largest *F* statistic, dashed vertical line is cue onset. Right: grand average topographies for each condition. B. Statistical analysis of the High>Low cue contrast in source-space found greater early activation of the ventral visual pathway (top: inferior temporal lobe) and late activation of visual, somatosensory and multimodal brain regions represented here in separate colours (for simplification, similar bilateral regions are coloured the same). C. Multivariate pattern analysis (MVPA) weights (top) and their projections (bottom). Maximal separation occurs between the two cue conditions at 485ms, at which time the topographic map (bottom) shows a posterior scalp distribution of activity that separates the two cues. In both time-course scatter plot, each point is a summarised weight or projection value (namely, the mean absolute value over the scalp and within a 10ms window). Y axis values are normalised to the % contribution, i.e. each data point is a percentage of the sum of all data points over time. D. Receiver Operating Characteristic (ROC) and MVPA statistics. The ROC illustrates the model sensitivity and specificity by plotting the true positive rate (sensitivity) as a function of false positive rate (1 - model specificity). The area under the curve (AUC) measures how well the model classifies the cue conditions (greater area means better classification).

**Figure 3:**
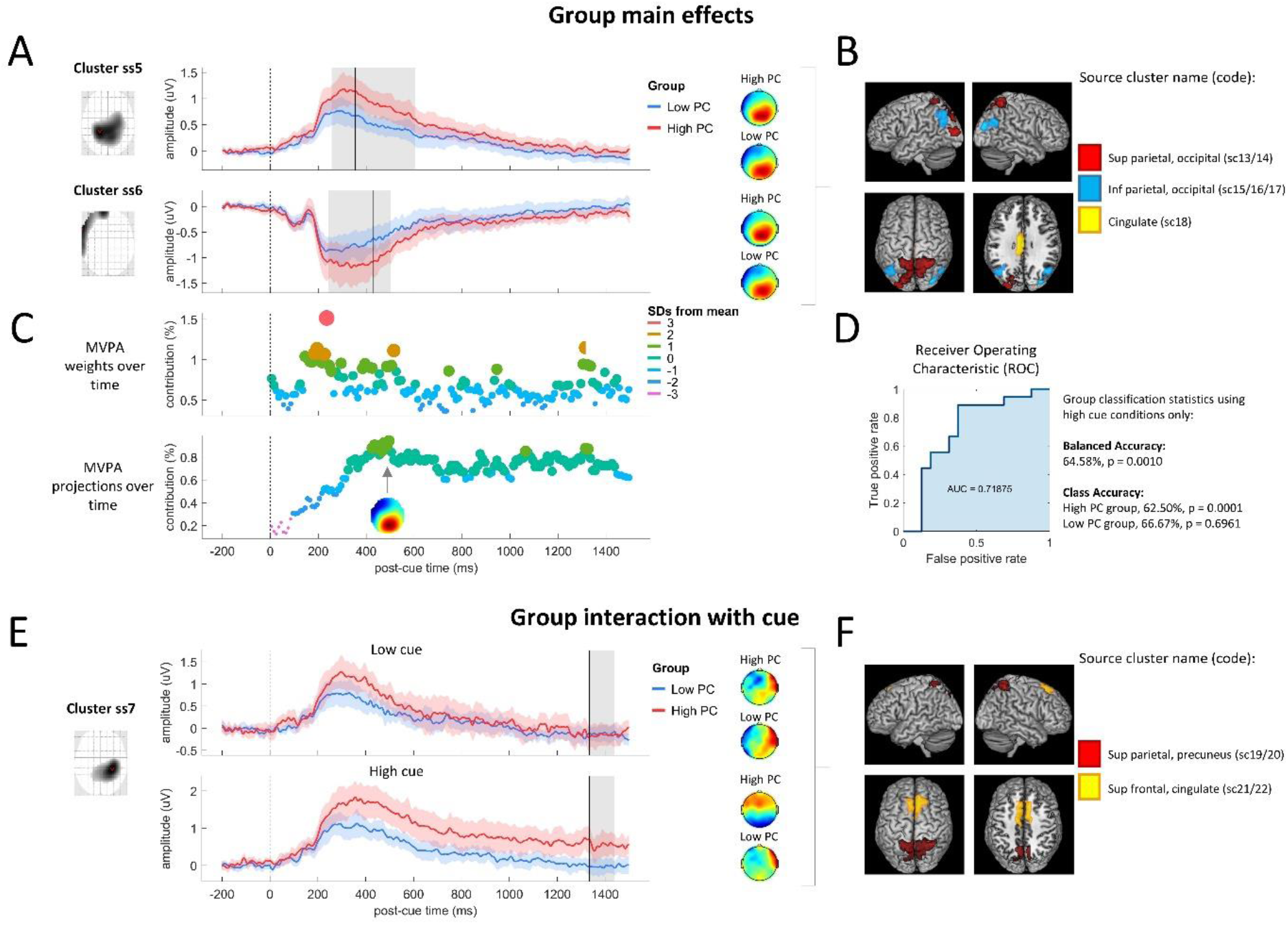
Group differences in cue processing (low PC: N=18, high PC: N=16). A. Univariate sensor analysis found two statistical clusters in the main effect of group. Clusters ss5 and ss6 are positive and negative polarities of the same ERP components during the time window of 242ms to 604ms. B. Source analysis found greater activation of occipital, parietal and cingulate cortex during cue processing in the contrast of High PC > Low PC group. C. Multivariate pattern analysis (MVPA) weights (top) and their projections (bottom) summarised within 10ms windows showing that neural activity maximally classifying the groups (at 495ms) occurs with a posterior scalp distribution. D. Receiver Operating Characteristic (ROC) and MVPA statistics for classification of group membership using the high cue condition only. AUC: Area under the ROC curve. E. Univariate sensor analysis of the interaction between group and cue type found a statistical cluster during the time window of 1334ms to 1440ms, resulting from a group difference in processing High cues but not Low cues. F. Source analysis found interaction effects in superior frontal-parietal cortex and cingulate cortex resulting from greater processing of High (vs. Low) cues in the High (vs. Low) PC group. See figure 2 legend for further details of the plots.

**Figure 4:**
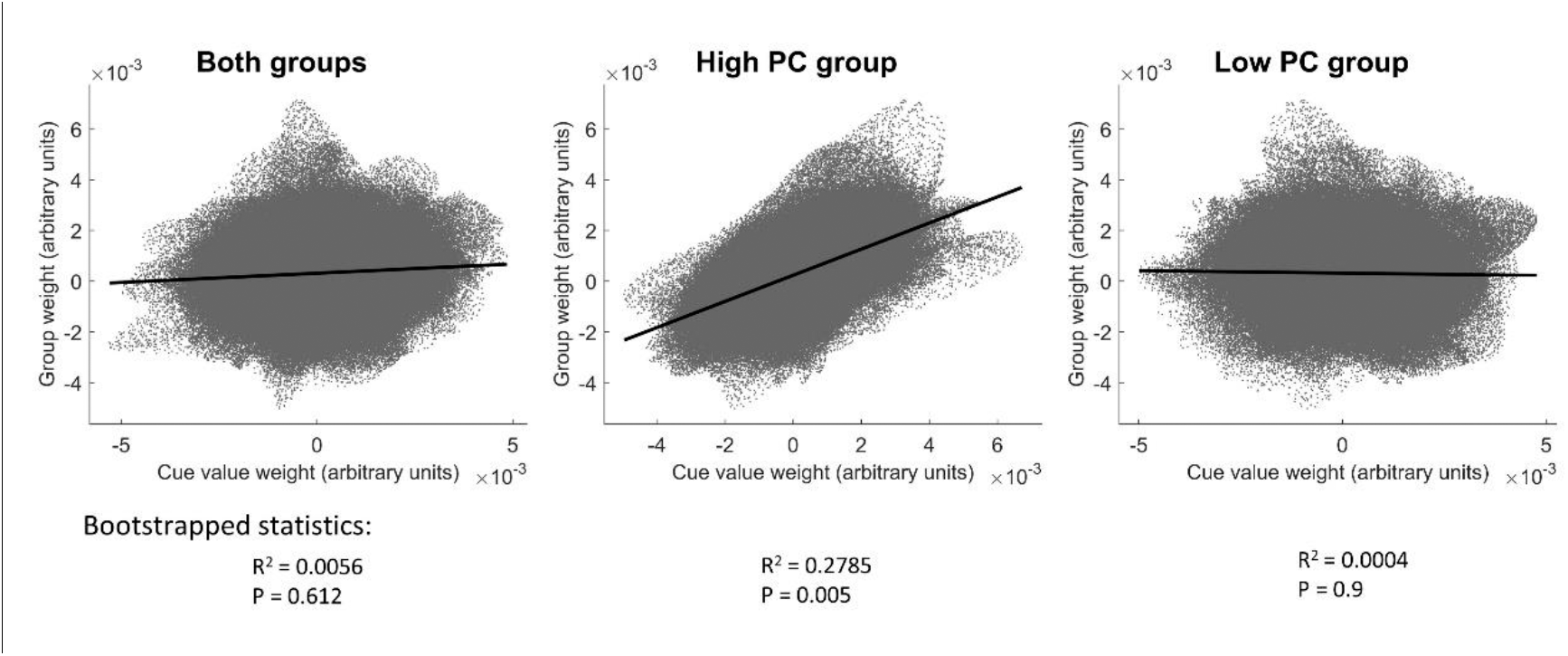
Predicting group classifier weights from cue classifier weights. Group classifier weights (y axis) are plotted against cue valuation weights (x axis). Cue classifier weights (x axis) are compared from three classification models: when classifying cues from both groups’ data (left) or from each group separately (middle and right).

In particular, univariate EEG sensor analysis on the within-subjects aversive valuation effect (figure 2A) identified spatiotemporal statistical clusters arising from increased positive and negative polarities co-occurring at similar latencies (N.B. greater neural activity / processing resources are indicated by absolute activity and not whether it is positive or negative; these polarities merely indicate opposite ends of an underlying source dipole). This occurred in two distinct latency ranges: an early latency range from 116ms to 152ms and a late latency range from 362ms to 1312ms post-cue (supplementary table 6). Source analysis over these time ranges revealed early latency sources in inferior temporal lobes, consistent with activation of the ventral visual pathway, while late latency sources localised to a broader range of regions including occipital, parietal, frontal, temporal and cingulate cortices (summarised in figure 2B and detailed in supplementary table 7).

Continuing our analysis of the within-subject aversive valuation effect, we used MVPA to train a classifier on the High vs. Low cue conditions, which provided a 69.12% balanced accuracy in classifying the two conditions (p = 0.001). We investigated the latencies of neural activity contributing to this classification, firstly with regard to raw MVPA weights (figure 2C), which provide information about spatiotemporal regions contributing the most to classification. In this analysis, there were moderately strong weight contributions (> 1 SD from the mean) across the full time range, with the strongest weight contribution at 455ms post-cue (> 2 SDs from the mean). However, because larger raw weights do not directly imply more class-specific information than lower weights, we transformed the classifier weights back into activation patterns (Haufe et al., 2014), providing an interpretable time-course (figure 2C, bottom). This shows latencies with the largest condition differences described by the classification, and produced a time-course distinct from the original ERP with a steady increase up to a maximum at 485ms. The topographic projections of the weights across the scalp at 485ms (figure 2C) showed a mid-posterior positivity and frontal negativity, reminiscent of (but not identical to) the ERP topography at this latency (figure 2A).

Having characterised the within-subject aversive valuation effect, we now turn our attention to group effects on neural activity in the 1500ms following the initial cue (figure 3). Univariate sensor analysis results show a main effect of group (high PC > low PC, across both cue conditions) at a mid-to-late latency range; two spatiotemporal sensor clusters were identified with the range of 242ms to 604ms, with greater activity in the high PC group (figure 3A). Topographic maps show that these two clusters correspond to concurrently occurring positive and negative polarities. Source analysis on this time range (figure 3B) locates the activity to superior and inferior parietal lobe and occipital lobe bilaterally, in addition to mid-cingulate cortex. The group effects therefore overlap both temporally (in the range 362ms to 502ms) and spatially (in the parietal lobes) with the aversive valuation effect (high vs. low cues)

Interaction *F* contrasts from the general linear model revealed a spatiotemporal cluster in the ultra-late latency range from 1334ms to 1440ms, characterised by greater group differences (high PC > low PC group) in the high cue compared to the low cue conditions (figure 3E). This latency window is considerably later than that found in the analysis of the main effect of group. Visually, as depicted in figure 3E, the interaction appears to arise from a group difference after low cues at mid-latencies (roughly, 300-400ms), but no group effect thereafter; whereas, after high cues, a similar group effect at mid-latencies persists into the ultra-late time window. (In the next section, we investigate whether this suggests more persistent expectancy processes that might carry over to affect nociceptive processing). We identified sources of these interaction effects as originating from superior parietal lobes, precuneus, superior frontal lobes (supplementary motor area) and mid-cingulate cortex (figure 3F).

MVPA results show general agreement with the above univariate results when assessing which spatiotemporal characteristics of the EEG data classify the two groups (figure 3C). Additionally, as a multivariate method, MVPA takes into account dependencies over time and space, and provides spatiotemporal patterns classifying the two groups that we later compare to the patterns classifying the two cue conditions. Firstly, two MVPA analyses were conducted to find out if successful group classification depends on what EEG data is used – here we look at results using data from high and low cue conditions separately. Interestingly, group was decoded from high cue conditions (64.58% balanced accuracy, p=0.0010, figure 3D) but not from low cue conditions (37.85% balanced accuracy, p=.9964, not shown as a figure). For the successful classification using high cue conditions, relatively early latencies (185ms to 245ms post-cue) contributed strongly to the weights (greater than 2 standard deviations from the mean weight contribution) with the greatest weight (>3SD) at 235ms post cue (figure 3C, top). However, as previously discussed, such weights might not be physiologically meaningful; for example they may reflect suppression of noise to improve the classification. Hence, we analysed the time-course of weight projections contributing to this classification (figure 3C, bottom) which showed a very similar temporal profile to the weight projections contributing to classification of the cue conditions (figure 2C), with the largest response at 495ms. The topography of the group weight projection at 495ms also showed a similar distribution to that classifying the cue conditions.

The above observations of a close spatiotemporal relationship between weight projections classifying groups on the one hand, and those classifying aversive value on the other, points to the possibility that *neural representations that successfully decode groups can be partially explained by representations of aversive value*. Indeed, further regression analyses support this view. Three regression analyses were conducted on MVPA raw weight matrices (figure 4). Firstly, we found that variance in the weight matrix that successfully decoded the groups was not explained by variance in the weight matrix decoding the cue conditions when both groups’ EEG data was used in the cue-condition classification analysis (R^2^ = 0.0056, p=0.612). However, as an alternative predictor variable, when we instead utilised the weight matrix from classifying cue-conditions using only the high PC group’s data, there was significant prediction of the variance in the group classification weights (R^2^ = 0.2785, p=0.005). In the third regression, the cue-classification weights from the low PC group did not predict the group classification weights (R^2^ = 0.0004, p=0.9). Hence, aversive value encoding as expressed specifically in the high PC group, but not in the low PC group, partially explains the pattern that decodes group membership.

### 3.4 Is pain catastrophizing characterised by prior pain expectancy effects on nociception?

We addressed this second question using the same methodology as in the previous section, except analyses were conducted on the post-stimulus Laser-Evoked Potential (LEP). However, as explicated in the following paragraphs, these analyses did not yield group differences.

As a preliminary step, we conducted univariate and multivariate analysis on the LEP to identify spatiotemporal representations of pain intensity (Contrast 2, figure 1C); this acted as validation of the analysis methods for the study as a whole, by enabling comparison of the latencies and sources of LEP components in this study to those found in previous research. These findings are in supplementary results and figure 5 A-D and are not referred to further here. In this current section, we instead focus on the investigation of the effects of prior pain expectation on LEP responses to medium intensity stimuli (Contrast 3 shown in figure 1C) and whether this effect was augmented in the high PC group.

**Figure 5:**
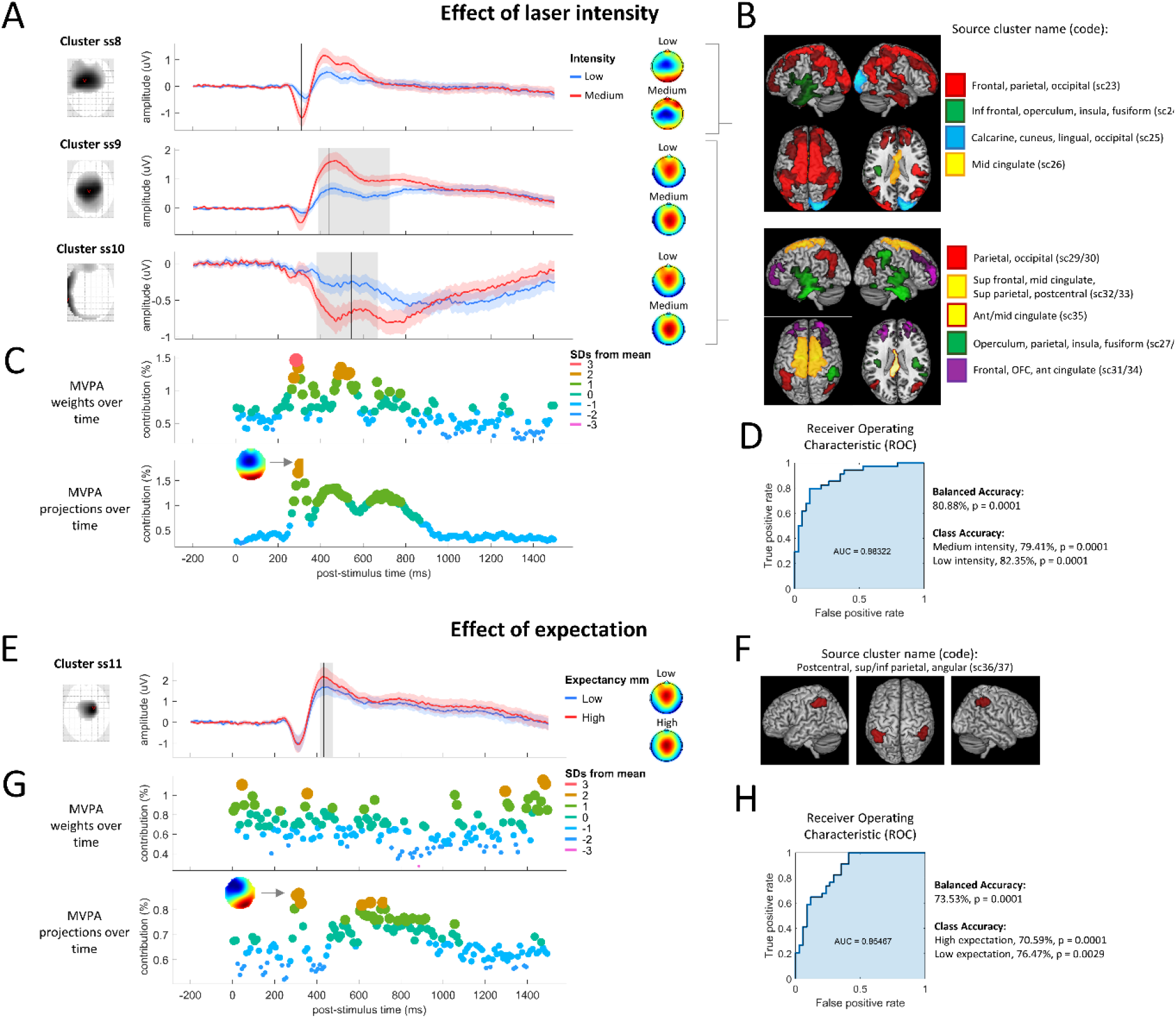
Effects of laser intensity (contrast 2, N=34) and prior expectation (Contrast 3, N=34) on nociceptive processing. A. Univariate sensor analysis found three statistical clusters from increasing laser intensity (medium>low), corresponding to components N2 (ss8, 308ms to 310ms) and P2/P3 (ss9, 388ms to 726ms; ss10, 382ms to 670ms). B. Source analysis found widespread cortical modulation in response to increases in pain intensity. C. Multivariate pattern analysis (MVPA) weights (top) and their projections (bottom) found that intensity was classified with the greatest contribution from neural activity during early latencies (N2 time window) and a fronto-central scalp distribution. D. Receiver Operating Characteristic (ROC) and MVPA statistics. AUC: Area under the ROC curve. D. Univariate sensor analysis found a single cluster from the prior pain expectancy effect, corresponding to component P2 (ss11, 416ms to 478ms). E. Source analysis found that prior pain expectancy increases post-laser stimulus activation of the inferior parietal lobes. F. Multivariate pattern analysis (MVPA) weights (top) and their projections (bottom) found that classification of the expectancy conditions involved neural activity patterns at both early latencies (N2 time window) and late latencies (P2/P3). H. Receiver Operating Characteristic (ROC) and MVPA statistics. AUC: Area under the ROC curve. See figure 2 legend for further details of the plots.

Univariate sensor analyses found neural activity related to the presentation of a prior high pain cue to be maximal at a latency of 434ms post-stimulus (cluster range: 416 to 478ms), consistent with the commonly observed P2 peak (figure 5E). At this latency, the P2 was maximal at electrode FCz. Source analysis revealed a contribution of bilateral inferior parietal cortex to this effect (figure 5F). Classification of the two conditions using MVPA provided 73.53% classification accuracy (p = 0.001). The latencies most greatly contributing to the weights (> 2 standard deviations from the mean) were far broader than that of the P2 peak, covering the full latency range of the window analysed. However, weight projections (figure 5G, bottom) identified more defined temporal regions at the latency of the N2 peak (maximal at 315ms) and at a later latency between the P2 and P3 peaks (605ms to 715ms). Topographic projections at 315ms were consistent with the commonly observed N2 peak of the LEP. The discrepancy between these latencies and those identified from the univariate results (i.e. P2 peak) highlights that univariate and multivariate analyses are sensitive to different signals: absolute mean activity within spatiotemporally restricted areas vs. relative patterns of activity over distributed regions respectively.

Despite significant univariate and multivariate EEG sensor differences between conditions, there was no evidence of a main effect of group or interaction between group and condition using mass univariate statistics. Likewise, MVPA was not able to obtain a statistically significant classification of the groups, either for spatiotemporal sensor data from the prior pain expectation condition (50%, p=0.9), nor using data from the low expectation condition (47.06%, p=0.9).

## 4 Discussion

In this paper we sought to address the general problem of a lack of viable biomarkers of the neurocognitive mechanisms influencing physical health outcomes. We specifically sought to identify neural representations of pain catastrophizing (PC) in healthy individuals using an experimental pain model. We found evidence in favour of a “magnification” hypothesis, namely augmented neural processing of cues predicting aversive (vs. non-aversive) outcomes, suggestive of magnified aversive valuation processes, which persisted for up to 1.5s after the aversive cue. MVPA provided moderately successful classification of high (relative to low) PC when applied specifically to processing of high aversive value cues, but not to the processing of low aversive cues, suggesting that neural representations of highly aversive cues best characterise pain catastrophizers. Indeed, this was further suggested by analyses showing that MVPA patterns from group classification were partially predicted by those patterns classifying aversive value in high pain catastrophizers specifically, suggesting an overlap in neural representations of aversive valuation and PC. This provides evidence supporting aversive valuation processes in the brain as potential biomarkers of cognitive/emotional processes related to PC.

On the other hand, we did not find evidence that expectancy effects on nociception and pain were more persistent in the high vs. low PC group after initially negative expectations were updated by contrary information indicating no threat of pain. We previously found that pain perception in healthy individuals undergoing a similar procedure were still influenced by expectations from the initial highly aversive cue, despite the presentation of a second cue to update expectation (the so-called ‘prior expectancy effect’ (Almarzouki et al., 2017)). However, in this study, although neural activity directly after aversive cues (prior to the second update cues) appeared to be more persistent over the 1.5s analysed in the high vs. low PC group, pointing to greater pain expectancy, none of the analyses (behavioural, univariate EEG or multivariate EEG) provided evidence that prior pain expectancy had a greater influence on pain and nociception in the high PC group. Overall, these findings suggest that anticipatory processes may be the most fruitful area for future investigation of biomarkers of PC mechanisms. In what follows, we discuss the nature of the biomarkers identified in this study and make comprehensive suggestions for future research.

By combining univariate and multivariate analysis approaches to the EEG data, we were able to both identify biomarkers (spatiotemporal patterns) classifying PC groups as well as gain insight into the timing and spatial localisation in the brain of these biomarkers. Identification of the latencies of cue-evoked responses that best differentiate high and low catastrophizers has two main benefits. Firstly, it provides links to previous research that has discovered specific task or other context-dependent modulation of neural activity at these latencies. This enables interpretation (although initially speculative) of the possible cognitive and neural mechanisms underlying group differences. Secondly, this facilitates the design of more nuanced experimental designs/tasks and EEG recording parameters that might better pinpoint the mechanisms of interest.

Specifically, we found that the topographies and timing of neural representations associated with PC (i.e. derived from transformed weight matrices from the MVPA analyses) are consistent with the commonly observed P3b (“endogenous” P3) and late-positivity waves of the visual-evoked potential, peaking in mid-posterior scalp regions at around 450-500ms post-cue. This positivity is commonly evoked by task-relevant stimuli and represents activity in multimodal networks thought to be involved with maintaining and updating representations of the task (Polich, 2009). Our source analysis found both aversive valuation and PC effects at this latency to originate from superior parietal and cingulate cortex. These regions may reflect one or a number of augmented cognitive processes in individuals with high PC. For example, a current neurobiological model of attention posits a dorsal frontoparietal network (including superior parietal lobule and dorsolateral prefrontal cortex (DLPFC)) as mediating top-down attention (Corbetta et al., 2008, 2002), a network that can be recruited via midcingulate cortex when signalling the need for greater cognitive control (Ridderinkhof et al., 2004). This is interesting in light of the observation that it is a distinct *ventral* frontoparietal network that primarily responds to salient stimuli such as pain (Downar et al., 2003) or its anticipation (Wiech et al., 2010). Greater dorsal frontoparietal responses in individuals with high PC therefore may therefore not necessarily indicate magnification of the salience of aversive stimuli, but rather a gain on the recruitment of subsequent cognitive control mechanisms. However, our analyses are not able to delineate how or whether these precise mechanisms contribute to PC. Important questions for the future are whether this source activity reflects greater excitatory or inhibitory activity in cortical neurons and whether increases in activity reflect changes in bottom-up or top-down streams of information processing.

The results highlight the utility of EEG for identifying temporally-defined neural patterns that could be the focus of further biomarker studies. However, despite the use of cross-validation to provide greater predictive validity to the results, there are clear limits on our ability to generalise the results of this study. Most importantly, the biomarkers were identified in (relatively young) healthy volunteers and so may not generalise to chronic pain patients. Furthermore, participants may not even be typical of the healthy population: respondents to the initial screening procedure were self-selected, with unknown factors underlying this; furthermore, approximately one third of screened participants were excluded from further participation due to being neither high nor low pain catastrophizers. There are also limitations on our ability to claim that differences in neural markers of cue/anticipation processing between groups are specific to the PC construct, as we were not able to measure and control for all extraneous variables. Of those we did measure, it would not be possible to distinguish states of PC in this study from states of anxiety (for which we also observed a group difference), given that the painful context of the experiment would be expected to induce anxiety in the high PC group. Furthermore, this type of study is not able to draw causal inferences between PC, consequent states such as anxiety, and the EEG outcomes; hence, we suggest that we have not identified specific markers of PC as such, but rather biomarkers of cognitive/emotional states related to PC.

We therefore view the results of the current study as providing an initial indication of neural targets, the robustness of which can be tested later as predictors of PC in chronic pain patients under a variety of contexts. These contexts might include alternative experimental designs that improve classification accuracy. Our design utilised very brief nociceptive stimulation of 100ms, which previous work has shown provides insights into the neural correlates of pain perception (Lee et al., 2009) and it’s modulation by expectation (Brown et al., 2008b). The brevity of the laser stimuli served two of our purposes well, which were to provide a relatively easy to tolerate stimulus that would be suitable for use in individuals who catastrophize about pain, allowing multiple trials of stimulation required to generate robust ERPs (Luck, 2005). However, despite these practical advantages, the use of brief nociceptive stimuli might not be an optimal method for inducing catastrophic cognitions. Support for our approach includes the moderate success we had in classifying high vs low PC groups using the neural pattern data and the fact that high PC participants rated themselves as expecting to be much more distressed by the laser stimuli than the low PC participants. However, future work could investigate whether pain catastrophizing is better classified by neural representations during more enduring and unpleasant tonic pain stimuli that provide a more realistic model of chronic pain. Future work could also investigate whether simpler expectation paradigms (i.e. those with single cues rather than changing cues) produce expectancy effects on nociception and pain that better differentiate high and low pain catastrophizers. This would help address one confound of our experimental design: the additional cues in conditions 3 and 4 compared to conditions 1 and 2 (see figure 1), mean that the “prior expectancy effect” contrast (contrast 3; results in figure 5E-H) may be contaminated by differences in perceived uncertainty or surprise. The current design does not facilitate differentiation of these uncertainty/surprise elements from the expectancy contrast, because of its focus on the *outcomes* of aversive learning rather than on the trial-by-trial learning *process* itself. On the other hand, certain types of more complex designs in combination with model-based inference of trial-by-trial hidden states (e.g. learning processes such as prediction errors and aversive value updating), which have previously been used to identify neural correlates of these hidden states (Seymour et al., 2004), may help to identify neural correlates of the learning processes themselves, namely trial-to-trial updating of expectation as distinct from uncertainty and surprise components. Such designs may also enable identification of anticipatory responses that are more specific to PC and to differentiate these from processes more specific to related cognitive/emotional states such as pain-related fear or generalised anxiety.

There are a number of further potential options for improving the identification of biomarkers of perseverative cognitions. Firstly, term “rumination” (Ottaviani et al., 2016) commonly refers to persistent, perseverative cognitions over longer timescales that we assessed here. Further research could aim to assess such persistent cognitions as well more naturalistically measure the dynamics of changes in pain expectancy (for example, during the course of a fluctuating tonic pain stimulus that mimics chronic pain symptoms). Secondly, regarding optimal neurophysiological measurements, neural representations of enduring cognitions may be better assessed by analysis of baseline or resting-state neural activity rather than using ERPs. Thirdly, a challenge for future studies is how to measure or influence rumination orthogonally to related cognitions such as magnification. The results of any such investigation will be highly dependent on how these variables are operationalised. Cross-sectional or longitudinal designs using questionnaire measures of trait rumination would require very large sample sizes to disentangle within or between-subject variance in rumination from that of related cognitive factors. For example, in our study, rumination was measured via one subscale of the PCS questionnaire, but the small sample size prohibited identifying variance specific to rumination and magnification. Alternatively, cognitive-behavioural interventions could be targeted to rumination, but such efforts are likely to produce knock-on effects to other cognitive variables.

Independently of issues of generalisation and optimal experimental design, the MVPA analyses presented here rely on certain assumptions and simplifications. Firstly, the study design leant itself to the use of binary classification; however, this approach assumes that PC is a unitary construct and that individuals in the high PC group differ from those in the low PC group according to homogeneous cognitive factors and corresponding neural representations; this is likely to be an over-simplification. Secondly, our analysis involved comparisons of weight matrices, but the interpretation of these values is complex, as weight values can vary either in relation to the signal of interest or in relation to their function in suppressing noise to improve prediction (Haufe et al., 2014). Hence raw weights do not have a simple neurophysiological interpretation. We therefore complimented raw weight information with a transformation of these weights as previously described (Haufe et al., 2014; Wardle et al., 2016) in order to enable interpretation of which physiological events contribute to the classification. However, other methods also exist for spatial and temporal localisation of neural signals contributing to multivariate classification (e.g. searchlight mapping (Etzel et al., 2013), sparse algorithms (Kampa et al., 2014), multiple kernel learning methods (Schrouff et al., 2018), etc.) and these methods merit exploration to assess consistency with our findings. Lastly, MVPA (like univariate analysis) benefits from the larger statistical power afforded by large sample sizes; the relative small sample size in this study was a limitation meaning that we had the power to detect only large effects.

## 5 Conclusion

The results of this study are consistent with the hypothesis that pain catastrophizing is characterised by magnification of aversive value encoding in cingulate and parietal cortices, possibly reflecting the initiation of cognitive control mechanisms. Our results provide an initial indication of neural targets that can be further tested as predictors of chronic pain and disability. Specifically, the results point to the importance of measuring anticipatory neural processes, as in our study these best differentiated high and low catastrophizing groups. This indicates that nociceptive responses may not provide sufficient information regarding mechanisms of interest to pain catastrophizing. We have highlighted the utility of EEG for identifying temporally-defined neural patterns that have potential as biomarkers of maladaptive cognition and we have pointed to the need for further research identifying optimal methodologies for biomarker development.

## 6 Acknowledgements

The authors declare no conflicts of interest. This research was funded by a PhD scholarship awarded to Abeer F. Almarzouki from the Government of Saudi Arabia.

## 7 Supplementary Methods

### 7.1 Latency windows used for EEG analysis

For both univariate and multivariate analyses of EEG data, analysis of cue processing was restricted to the first 1500ms following the first cue rather than utilising the whole window up until the laser stimulus. This served two purposes: (1) for MVPA, reducing the feature space as far as possible to minimise over-fitting, (2) removing ERP signals at around the time of 2000ms after each cue, at which time the offset of the visual cues occurred, which generated small additional visual-evoked responses that were not of interest in the analysis. The first 1500ms post-cue is likely sufficient to include both conventional visual evoked responses to the cues and subsequent anticipatory responses prior to the laser stimuli, which can be assumed to change as a matter of degree rather than kind from that point on. This is suggested by our previous work showing mid-range anticipatory responses having the same character as that of late-range (immediately pre-laser) responses (Brown et al., 2008a).

### 7.2 EEG sensor data: Mass univariate analysis (MUA)

Sensor analyses were conducted by converting sensor-by-time EEG data to three-dimensional images using the Statistical Parametric Mapping software (SPM12, www.fil.ion.ucl.ac.uk/spm) (Litvak and Friston, 2008), and subjecting these to mass univariate analysis (in SPM12) with correction for multiple comparisons using random field theory. This is the standard approach to EEG analysis implemented in SPM12 (Litvak et al., 2011). This analysis deals with the multiple comparisons problem (in this case, statistical inference over many peri-stimulus time points and many electrodes) in a way that does not require narrowing down the search space by making assumptions as to the precise timing or topographic location of physiologically important events. More precisely, random field theory (RFT) is used to make inferences over space and time while adjusting *p*-values in a way that takes into account the non-independence of neighbouring sensors and time-points (Litvak et al., 2011). Applying to smooth data, the RFT adjustment is more sensitive than a Bonferroni correction. Hence, after data conversion from EEGLAB to SPM format, for each participant, experiment and digit, SPM EEG sensor data were transformed to 3-D Scalp [x, y] × Time [z] images and smoothed in the spatial dimension to 12mm full-width half-maximum (FWHM).

General Linear Models (GLMs) were estimated at the group level consisting of the between-subject factors Subject and Group (High PC, Low PC), the within-subject effect of interest (either aversive valuation effect, pain intensity effect or prior expectancy effect in different models) and a further within-subject factor of Block. The Block variable was a nuisance variable included to account for data variability related to the different tasks in blocks 1 and 2. More specifically, the GLM model of the cue valuation effect (High cue > Low cue) included EEG data from the time window of 0ms to 1500ms after High cues (conditions 3 and 4) and contrasted to the same time window after Low cues (conditions 1 and 2; see figure 1). The pain intensity and prior expectancy effects on nociceptive processing were both modelled using the time window of 0ms to 1500ms post-laser stimulus. In both cases, EEG sensor images were first baseline corrected to the 500ms preceding the window-of-interest (relatively long 500ms baselines were used so that resulting ERP estimates were less susceptible to transient baseline noise). For each GLM, three F-contrasts were relevant: (1) the main within-subjects effect of interest (aversive valuation, pain intensity or prior expectancy effect), (2) the main effect of Group (between-subjects factor of interest), (2) the interaction of these between and within-subjects factors.

For any statistical models in which there was an imbalance in the number of trials between conditions in the design (for example, in contrast 2 between medium and low intensity laser stimuli), trials were randomly sampled (once only per subject/condition) from the condition with the larger number of trials to match the number in the conditions with the smaller number. This balancing was also critically important for subsequent MVPA analyses to ensure validity of the results.

A final consideration is as to whether to control for extraneous variables such as trait anxiety (which may confound the results owing to overlap with the construct of PC) in the statistical analyses. This was not done for the following reasons. Firstly, regression with multiple predictors requires larger sample sizes than regression with single predictors, which the study was not designed to accommodate; results would therefore be at greater risk type II error rates. Secondly, the use of multiple regression to establish incremental validity is associated with extremely high type I error rates (see Westfall and Yarkoni, 2016 for a detailed discussion). This means that conclusions in the literature that one construct contributes incrementally to an outcome, or that two constructs are theoretically distinct, are often unwarranted, although this depends on the reliability of the measurement. In particular, addressing questions of incremental validity is especially problematic when the nuisance covariate (e.g. trait anxiety scores on the STAI) is a noisy proxy for the underlying latent variable (e.g. trait anxiety), which is commonly a problem for self-report measures such as questionnaires.

### 7.3 EEG source analysis

Further analysis aimed to identify sources of the ERPs during those time windows that showed statistically significant effects in the sensor analysis. Canonical sensor locations were coregistered with the canonical head model in SPM12. Lead field computation used a boundary element model (Litvak et al., 2011). The Bayesian source reconstruction method in SPM12 was used with Multiple Spare Priors (Friston et al., 2008) to estimate sources across the temporal window-of-interest (the same windows as described above for sensor analysis). Subsequently, source activity was averaged in a series of smaller time clusters corresponding to the statistically significant effects from the sensor analyses. Using *F* contrasts, significant differences were identified in source space and reported significant at a cluster-level significance of *p* (FWE) < 0.05 when considering statistical maps thresholded at *p* < 0.001.

### 7.4 EEG sensor data: multivariate pattern analysis (MVPA)

MVPA involved learning classifiers on sensor images at the 2^nd^ (group) level using the Pattern Recognition for Neuroimaging Toolbox (PRoNTo) (Schrouff et al., 2013b). Classifiers are trained to identify patterns differentiating between the two levels of each within-subject factor of interest (cue type and prior expectation) and between groups at each time window of interest. The feature spaces used were the same three-dimensional spatiotemporal sensor images as used for MUA; when combined over subjects and conditions, these formed the feature vector for the Group or Condition targets that were fed into the classifier.

Regarding the classification algorithm and normalisation procedures, we used recommendations from a recent study of ERP biomarkers in Schizophrenia (Taylor et al., 2017). We used a Gaussian process classifier (GPC) (Rasmussen and Williams, 2004). GPC uses Bayesian modelling to estimate the likelihood that a test sample belongs to a particular class, by using the covariance structure of the data to make predictions and assign a class label. Performance of the classifier is assessed against the true target assignments. To avoid the problem of over-fitting and to improve generalisation of the results, parameter estimation is regularised and the performance of models were tested using a leave-one-subject-out-per-group cross-validation scheme, in which the classifiers were trained on data from all subjects bar one from each group, and tested on the excluded subjects. Across a number of ‘folds’, subjects were iteratively assigned for testing the model until all had been used once. During cross-validation, mean-centering was applied. Statistical significance of classification accuracy was assessed using permutation tests, involving retraining each model 10,000 times with randomised target labels.

### 7.5 MVPA weight matrix analyses

Analysis was conducted on the spatiotemporal weight matrices outputted from MVPA. Firstly, we identified temporal windows that contributed the most to the classification for each model, providing a supplement to univariate analysis of the temporal localisation of each effect. Because it is not meaningful to threshold the obtained weight map, we used *a posteriori* weight summarization (Schrouff et al., 2013a) in which local averages of the weights are obtained. In our case, local averages were calculated over the two spatial dimensions of the image (i.e. the scalp map) within consecutive non-overlapping 10ms time windows. Furthermore, because larger raw weights do not directly imply more class-specific information than lower weights, we used a solution introduced by Haufe et al. (Haufe et al., 2014) (previously applied to MEG decoding by Wardle et al. (Wardle et al., 2016)) which involves transforming the classifier weights back into activation patterns, providing an interpretable time-course. This involves multiplication of the weight matrices with the covariance in the EEG data used to derive those weights.

We also sought to identify the existence of shared representations contributing to both group and condition classifications. We quantified the degree of representational similarity (Kriegeskorte, 2008) by regressing vectorised spatiotemporal raw weight matrices from the classification of groups (pain catastrophizing effect) on those classifying the cue types (aversive valuation effect). *P*-values were calculated from bootstrap tests in which regressions were run for 1000 permuted classifications and their resulting weight matrices (this was kept to 1000 due to the computational load of outputting weight matrices using PRoNTo).

## 8 Supplementary Results

### 8.1 Participant numbers

163 participants (53 high PC, 65 low PC and 45 in-between) completed the PCS screening procedure and the high and low PC individuals were invited to participate in the EEG experiment. Of these, 36 high PC and 30 low PC attended the study visit, while the remainder did not accept the invitation. Upon repeat assessment using the PCS during the study visit, 13 high PC and 6 low PC no longer fell in the upper/lower quartile on the PCS respectively and were excluded (this was expected and consistent with previous findings in which the test-retest reliability of the PCS was been found to be 0.73 in a chronic pain population (Lamé et al., 2008)). Three further volunteers from the high PC group withdrew from the study due to discomfort from the laser stimulation or application of the EEG cap, while two participants were excluded from the low PC group because they did not find the stimulus painful. A further 3 participants’ data from each group was not included in the final analyses due to excessive artefact in the EEG data that could not be removed. One participant in each group was excluded from analysis due to failing the manipulation check, having reported not attending to the visual cues.

### 8.2 Sensor and source analysis of laser intensity coding

Initial validation of the univariate sensor and source analysis was conducted on the time range of the LEP by investigating the main effect of laser stimulus intensity (Contrast 2, see figure 1C for conditions included). The temporal and spatial effects of intensity were consistent with the findings of previous research. Intensity increased LEP amplitudes at latencies consistent with the commonly observed N2 and P2 peaks (figure 5A) at 308ms to 310ms and 382ms to 726ms respectively (supplementary table 6). Source results from the intensity contrast at each of these mid and late latencies were very similar (figure 5B and supplementary table 7), showing intensity modulation of widespread cortical regions. Importantly, these included commonly activated regions of the “pain matrix”, namely the insula, fronto-parietal operculum, primary somatosensory cortex, mid-cingulate cortex, plus a broader range of multimodal regions in frontal, parietal and temporal lobes. Classification of the two intensity conditions was conducted using MVPA, providing a strong classification accuracy at 80.88% (p = 0.0001). The latencies of the raw weights and transformed weights most contributing to the classification are consistent with the latencies of the N2, P2 and P3 components of the LEP, with a greater weighting towards the N2 latency range (maximal raw weight: 285ms; maximal transformed weight projection: 305ms).

## 9 Supplementary Tables

### 9.1 Supplementary Table 1: Participant Data

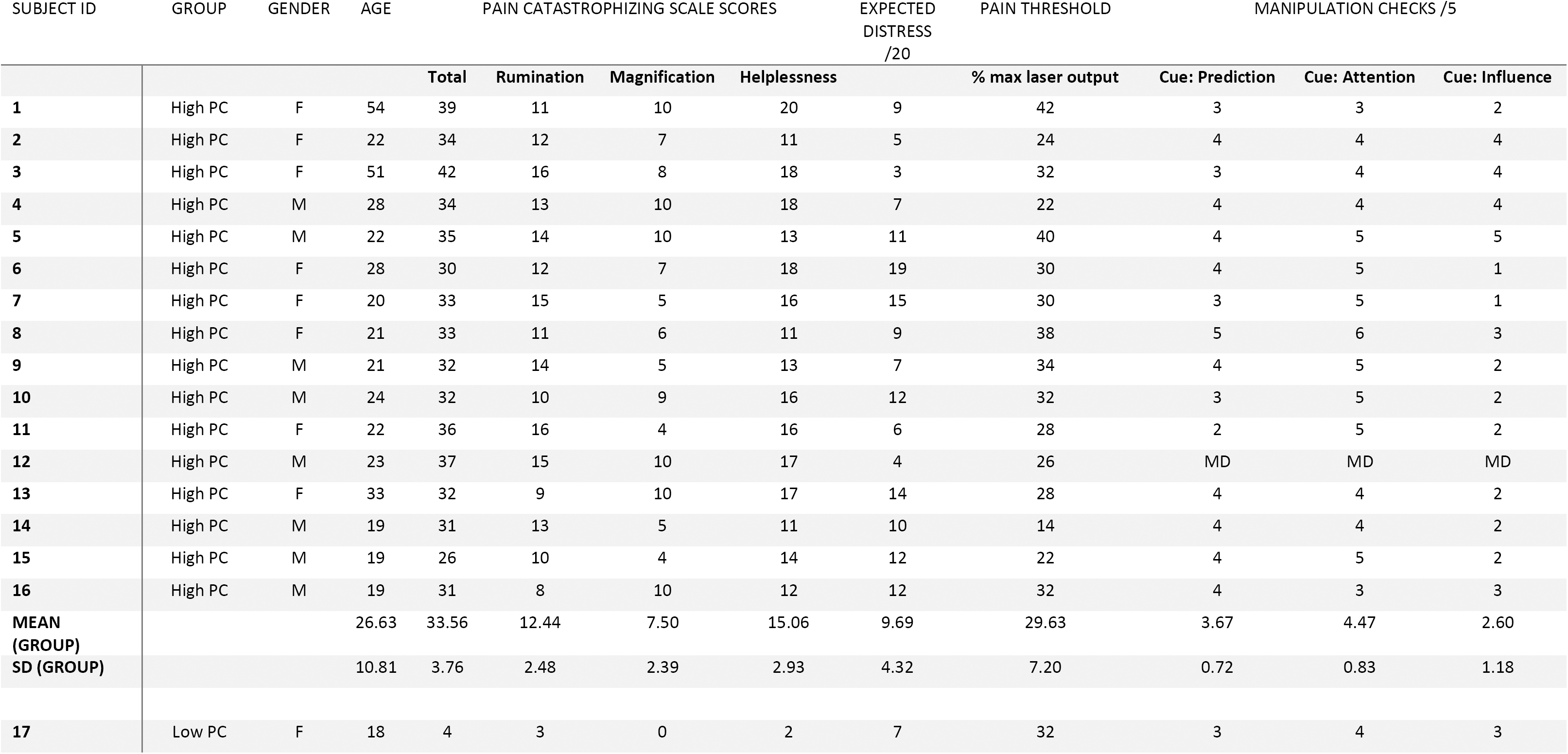
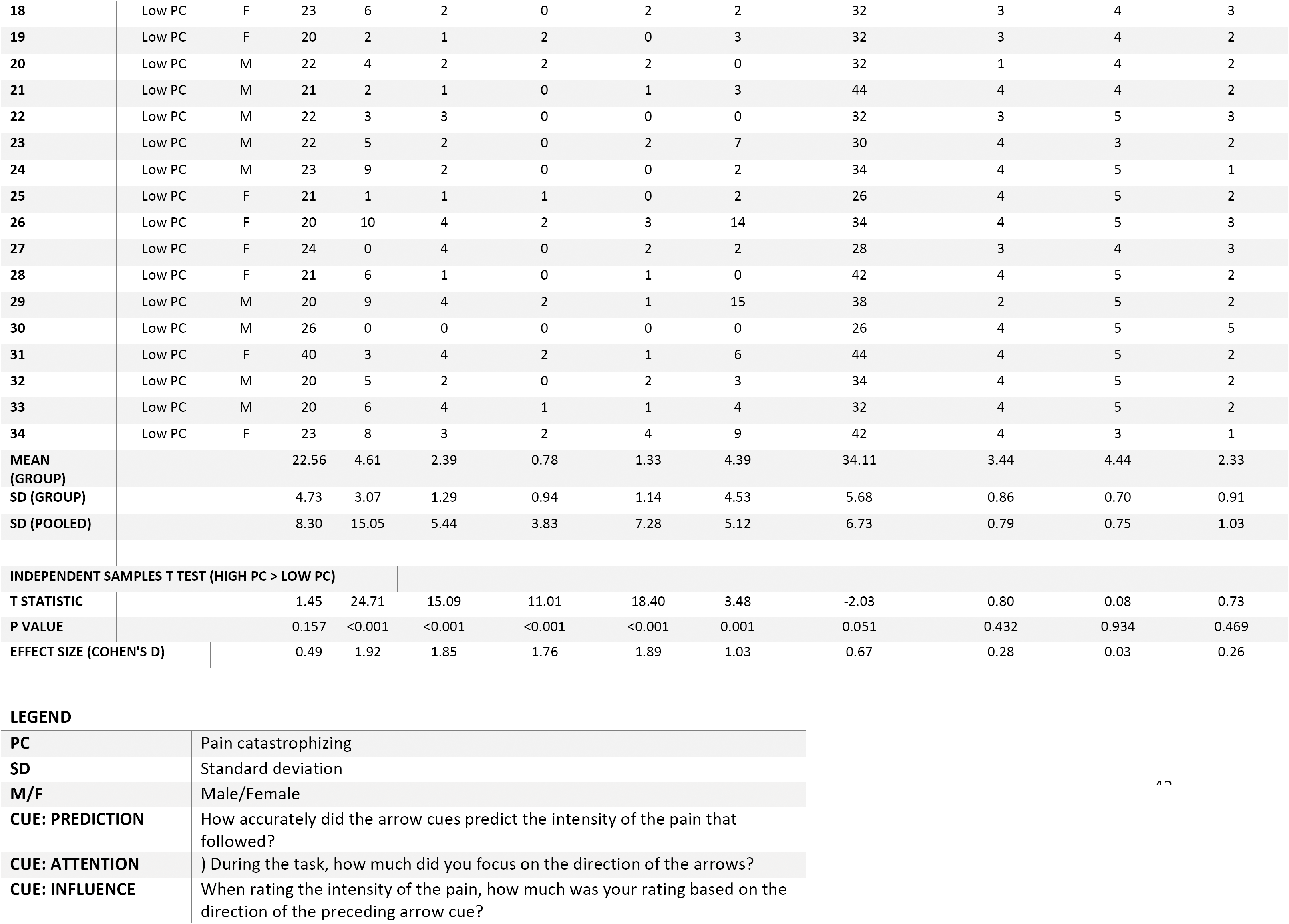

| SUBJECT ID | GROUP | GENDER | AGE | PAIN CATASTROPHIZING SCALE SCORES |  |  |  | EXPECTED<br>DISTRESS<br>/20 | PAIN THRESHOLD | MANIPULATION CHECKS /5 |  |  |
| --- | --- | --- | --- | --- | --- | --- | --- | --- | --- | --- | --- | --- |
|  |  |  |  | Total | Rumination | Magnification | Helplessness |  | % max laser output | Cue: Prediction | Cue: Attention | Cue: Influence |
| 1 | High PC | F | 54 | 39 | 11 | 10 | 20 | 9 | 42 | 3 | 3 | 2 |
| 2 | High PC | F | 22 | 34 | 12 | 7 | 11 | 5 | 24 | 4 | 4 | 4 |
| 3 | High PC | F | 51 | 42 | 16 | 8 | 18 | 3 | 32 | 3 | 4 | 4 |
| 4 | High PC | M | 28 | 34 | 13 | 10 | 18 | 7 | 22 | 4 | 4 | 4 |
| 5 | High PC | M | 22 | 35 | 14 | 10 | 13 | 11 | 40 | 4 | 5 | 5 |
| 6 | High PC | F | 28 | 30 | 12 | 7 | 18 | 19 | 30 | 4 | 5 | 1 |
| 7 | High PC | F | 20 | 33 | 15 | 5 | 16 | 15 | 30 | 3 | 5 | 1 |
| 8 | High PC | F | 21 | 33 | 11 | 6 | 11 | 9 | 38 | 5 | 6 | 3 |
| 9 | High PC | M | 21 | 32 | 14 | 5 | 13 | 7 | 34 | 4 | 5 | 2 |
| 10 | High PC | M | 24 | 32 | 10 | 9 | 16 | 12 | 32 | 3 | 5 | 2 |
| 11 | High PC | F | 22 | 36 | 16 | 4 | 16 | 6 | 28 | 2 | 5 | 2 |
| 12 | High PC | M | 23 | 37 | 15 | 10 | 17 | 4 | 26 | MD | MD | MD |
| 13 | High PC | F | 33 | 32 | 9 | 10 | 17 | 14 | 28 | 4 | 4 | 2 |
| 14 | High PC | M | 19 | 31 | 13 | 5 | 11 | 10 | 14 | 4 | 4 | 2 |
| 15 | High PC | M | 19 | 26 | 10 | 4 | 14 | 12 | 22 | 4 | 5 | 2 |
| 16 | High PC | M | 19 | 31 | 8 | 10 | 12 | 12 | 32 | 4 | 3 | 3 |
| MEAN<br>(GROUP) |  |  | 26.63 | 33.56 | 12.44 | 7.50 | 15.06 | 9.69 | 29.63 | 3.67 | 4.47 | 2.60 |
| SD (GROUP) |  |  | 10.81 | 3.76 | 2.48 | 2.39 | 2.93 | 4.32 | 7.20 | 0.72 | 0.83 | 1.18 |
| 17 | Low PC | F | 18 | 4 | 3 | 0 | 2 | 7 | 32 | 3 | 4 | 3 |
| 18 | Low PC | F | 23 | 6 | 2 | 0 | 2 | 2 | 32 | 3 | 4 | 3 |
| 19 | Low PC | F | 20 | 2 | 1 | 2 | 0 | 3 | 32 | 3 | 4 | 2 |
| 20 | Low PC | M | 22 | 4 | 2 | 2 | 2 | 0 | 32 | 1 | 4 | 2 |
| 21 | Low PC | M | 21 | 2 | 1 | 0 | 1 | 3 | 44 | 4 | 4 | 2 |
| 22 | Low PC | M | 22 | 3 | 3 | 0 | 0 | 0 | 32 | 3 | 5 | 3 |
| 23 | Low PC | M | 22 | 5 | 2 | 0 | 2 | 7 | 30 | 4 | 3 | 2 |
| 24 | Low PC | M | 23 | 9 | 2 | 0 | 0 | 2 | 34 | 4 | 5 | 1 |
| 25 | Low PC | F | 21 | 1 | 1 | 1 | 0 | 2 | 26 | 4 | 5 | 2 |
| 26 | Low PC | F | 20 | 10 | 4 | 2 | 3 | 14 | 34 | 4 | 5 | 3 |
| 27 | Low PC | F | 24 | 0 | 4 | 0 | 2 | 2 | 28 | 3 | 4 | 3 |
| 28 | Low PC | F | 21 | 6 | 1 | 0 | 1 | 0 | 42 | 4 | 5 | 2 |
| 29 | Low PC | M | 20 | 9 | 4 | 2 | 1 | 15 | 38 | 2 | 5 | 2 |
| 30 | Low PC | M | 26 | 0 | 0 | 0 | 0 | 0 | 26 | 4 | 5 | 5 |
| 31 | Low PC | F | 40 | 3 | 4 | 2 | 1 | 6 | 44 | 4 | 5 | 2 |
| 32 | Low PC | M | 20 | 5 | 2 | 0 | 2 | 3 | 34 | 4 | 5 | 2 |
| 33 | Low PC | M | 20 | 6 | 4 | 1 | 1 | 4 | 32 | 4 | 5 | 2 |
| 34 | Low PC | F | 23 | 8 | 3 | 2 | 4 | 9 | 42 | 4 | 3 | 1 |
| MEAN (GROUP) |  |  | 22.56 | 4.61 | 2.39 | 0.78 | 1.33 | 4.39 | 34.11 | 3.44 | 4.44 | 2.33 |
| SD (GROUP) |  |  | 4.73 | 3.07 | 1.29 | 0.94 | 1.14 | 4.53 | 5.68 | 0.86 | 0.70 | 0.91 |
| SD (POOLED) |  |  | 8.30 | 15.05 | 5.44 | 3.83 | 7.28 | 5.12 | 6.73 | 0.79 | 0.75 | 1.03 |
| INDEPENDENT SAMPLES T TEST (HIGH PC > LOW PC) |  |  |  |  |  |  |  |  |  |  |  |  |
| T STATISTIC |  |  | 1.45 | 24.71 | 15.09 | 11.01 | 18.40 | 3.48 | -2.03 | 0.80 | 0.08 | 0.73 |
| P VALUE |  |  | 0.157 | <0.001 | <0.001 | <0.001 | <0.001 | 0.001 | 0.051 | 0.432 | 0.934 | 0.469 |
| EFFECT SIZE (COHEN'S D) |  |  | 0.49 | 1.92 | 1.85 | 1.76 | 1.89 | 1.03 | 0.67 | 0.28 | 0.03 | 0.26 |

LEGEND
|  |  |
| --- | --- |
| PC | Pain catastrophizing |
| **SD** | Standard deviation |
| **M/F** | Male/Female |
| **CUE: PREDICTION** | How accurately did the arrow cues predict the intensity of the pain that followed? |
| **CUE: ATTENTION** | ) During the task, how much did you focus on the direction of the arrows? |
| **CUE: INFLUENCE** | When rating the intensity of the pain, how much was your rating based on the direction of the preceding arrow cue? |

### 9.2 Supplementary Table 2: Participant Data (EXTRANEOUS VARIABLES)

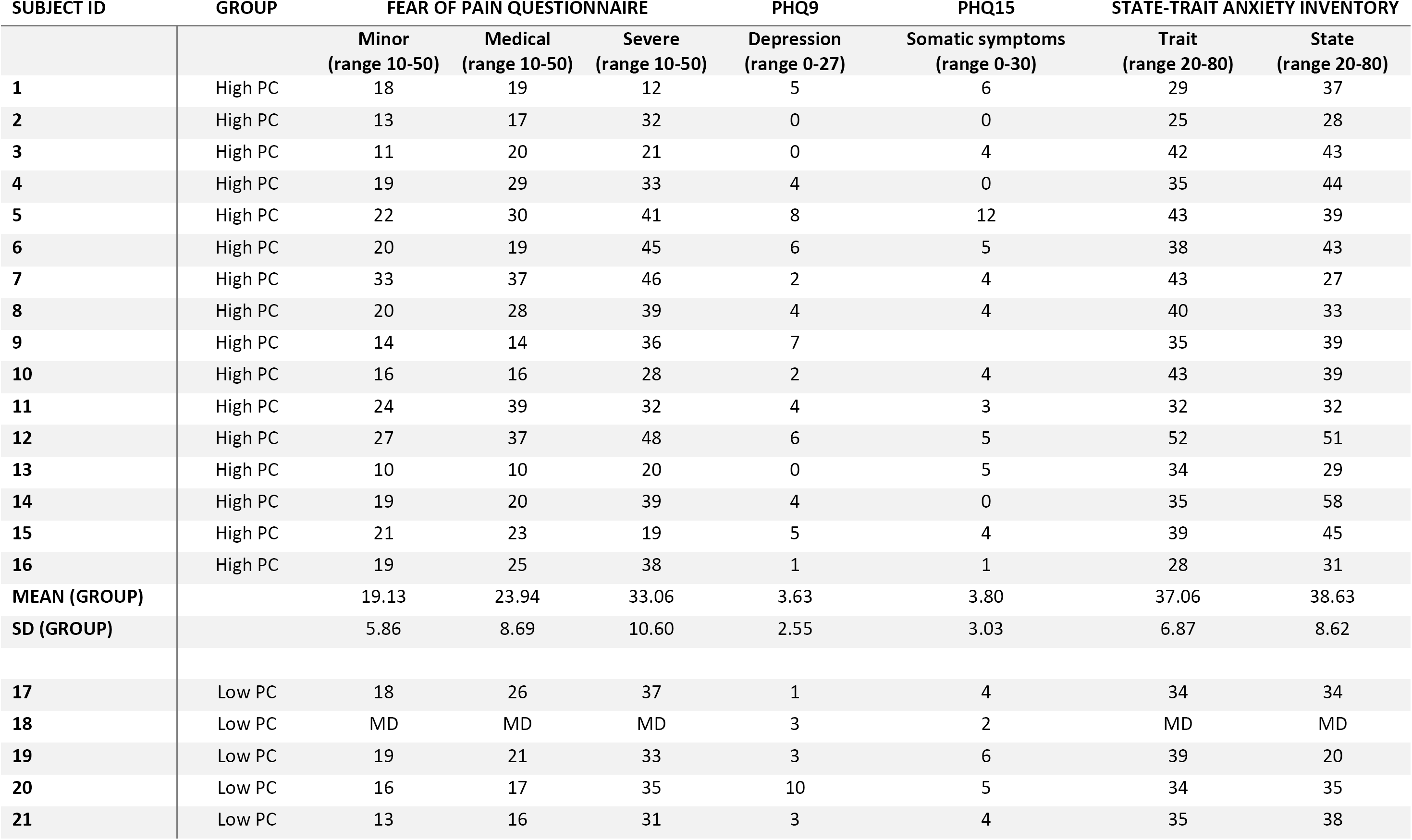
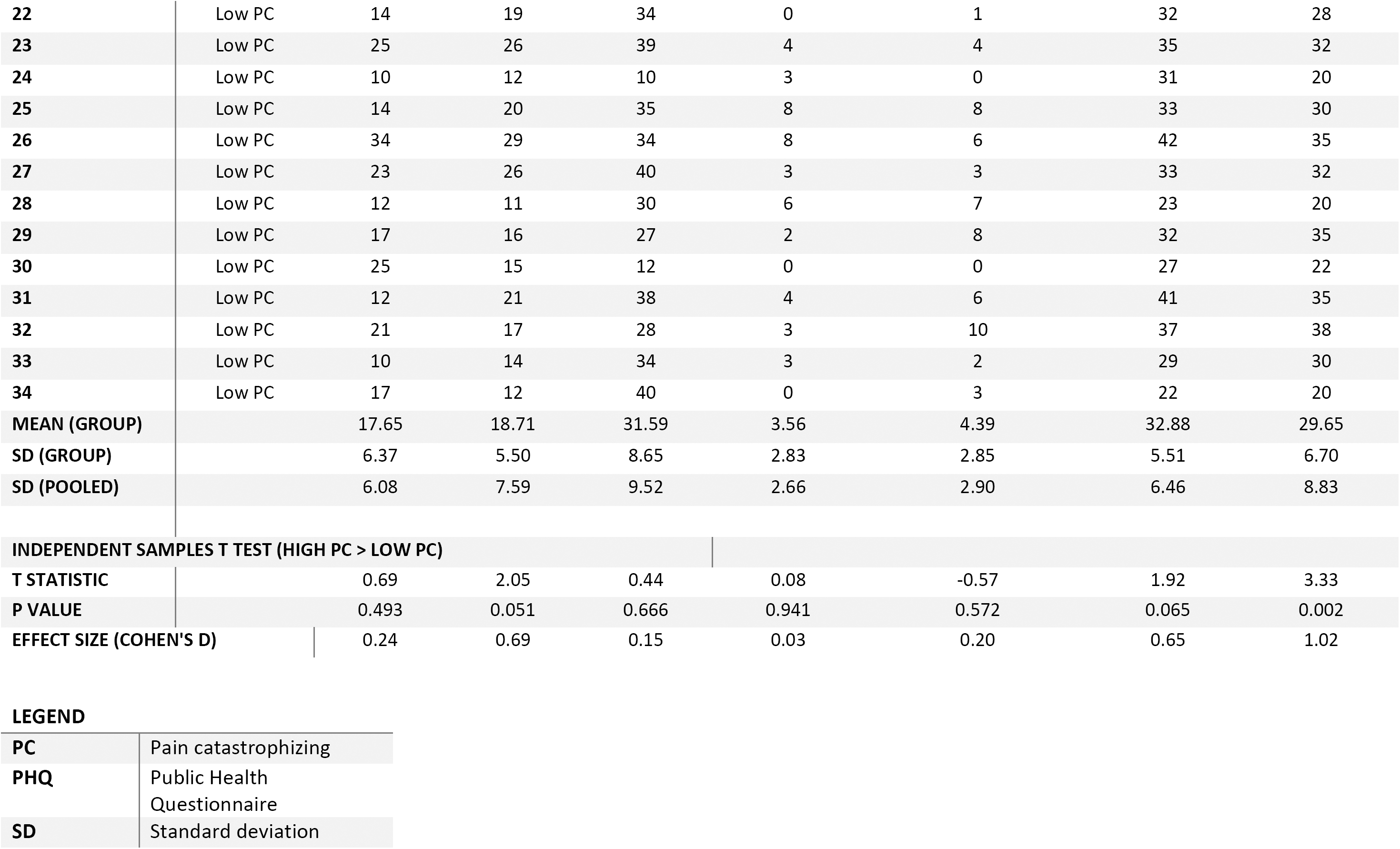

| SUBJECT ID | GROUP | FEAR OF PAIN QUESTIONNAIRE |  |  | PHQ9 | PHQ15 | STATE-TRAIT ANXIETY INVENTORY |  |
| --- | --- | --- | --- | --- | --- | --- | --- | --- |
|  |  | Minor<br>(range 10-50) | Medical<br>(range 10-50) | Severe<br>(range 10-50) | Depression<br>(range 0-27) | Somatic symptoms<br>(range 0-30) | Trait<br>(range 20-80) | State<br>(range 20-80) |
| 1 | High PC | 18 | 19 | 12 | 5 | 6 | 29 | 37 |
| 2 | High PC | 13 | 17 | 32 | 0 | 0 | 25 | 28 |
| 3 | High PC | 11 | 20 | 21 | 0 | 4 | 42 | 43 |
| 4 | High PC | 19 | 29 | 33 | 4 | 0 | 35 | 44 |
| 5 | High PC | 22 | 30 | 41 | 8 | 12 | 43 | 39 |
| 6 | High PC | 20 | 19 | 45 | 6 | 5 | 38 | 43 |
| 7 | High PC | 33 | 37 | 46 | 2 | 4 | 43 | 27 |
| 8 | High PC | 20 | 28 | 39 | 4 | 4 | 40 | 33 |
| 9 | High PC | 14 | 14 | 36 | 7 |  | 35 | 39 |
| 10 | High PC | 16 | 16 | 28 | 2 | 4 | 43 | 39 |
| 11 | High PC | 24 | 39 | 32 | 4 | 3 | 32 | 32 |
| 12 | High PC | 27 | 37 | 48 | 6 | 5 | 52 | 51 |
| 13 | High PC | 10 | 10 | 20 | 0 | 5 | 34 | 29 |
| 14 | High PC | 19 | 20 | 39 | 4 | 0 | 35 | 58 |
| 15 | High PC | 21 | 23 | 19 | 5 | 4 | 39 | 45 |
| 16 | High PC | 19 | 25 | 38 | 1 | 1 | 28 | 31 |
| MEAN (GROUP) |  | 19.13 | 23.94 | 33.06 | 3.63 | 3.80 | 37.06 | 38.63 |
| SD (GROUP) |  | 5.86 | 8.69 | 10.60 | 2.55 | 3.03 | 6.87 | 8.62 |
| 17 | Low PC | 18 | 26 | 37 | 1 | 4 | 34 | 34 |
| 18 | Low PC | MD | MD | MD | 3 | 2 | MD | MD |
| 19 | Low PC | 19 | 21 | 33 | 3 | 6 | 39 | 20 |
| 20 | Low PC | 16 | 17 | 35 | 10 | 5 | 34 | 35 |
| 21 | Low PC | 13 | 16 | 31 | 3 | 4 | 35 | 38 |
| **22** | Low PC | 14 | 19 | 34 | 0 | 1 | 32 | 28 |
| **23** | Low PC | 25 | 26 | 39 | 4 | 4 | 35 | 32 |
| **24** | Low PC | 10 | 12 | 10 | 3 | 0 | 31 | 20 |
| **25** | Low PC | 14 | 20 | 35 | 8 | 8 | 33 | 30 |
| **26** | Low PC | 34 | 29 | 34 | 8 | 6 | 42 | 35 |
| **27** | Low PC | 23 | 26 | 40 | 3 | 3 | 33 | 32 |
| **28** | Low PC | 12 | 11 | 30 | 6 | 7 | 23 | 20 |
| **29** | Low PC | 17 | 16 | 27 | 2 | 8 | 32 | 35 |
| **30** | Low PC | 25 | 15 | 12 | 0 | 0 | 27 | 22 |
| **31** | Low PC | 12 | 21 | 38 | 4 | 6 | 41 | 35 |
| **32** | Low PC | 21 | 17 | 28 | 3 | 10 | 37 | 38 |
| **33** | Low PC | 10 | 14 | 34 | 3 | 2 | 29 | 30 |
| **34** | Low PC | 17 | 12 | 40 | 0 | 3 | 22 | 20 |
| **MEAN (GROUP)** |  | 17.65 | 18.71 | 31.59 | 3.56 | 4.39 | 32.88 | 29.65 |
| **SD (GROUP)** |  | 6.37 | 5.50 | 8.65 | 2.83 | 2.85 | 5.51 | 6.70 |
| **SD (POOLED)** |  | 6.08 | 7.59 | 9.52 | 2.66 | 2.90 | 6.46 | 8.83 |
| **INDEPENDENT SAMPLES T TEST (HIGH PC > LOW PC)** |  |  |  |  |  |  |  |  |
| **T STATISTIC** |  | 0.69 | 2.05 | 0.44 | 0.08 | -0.57 | 1.92 | 3.33 |
| **P VALUE** |  | 0.493 | 0.051 | 0.666 | 0.941 | 0.572 | 0.065 | 0.002 |
| **EFFECT SIZE (COHEN'S D)** |  | 0.24 | 0.69 | 0.15 | 0.03 | 0.20 | 0.65 | 1.02 |

LEGEND
|  |  |
| --- | --- |
| **PC** | Pain catastrophizing |
| **PHQ** | Public Health<br>Questionnaire |
| **SD** | Standard deviation |

### 9.3 Supplementary Table 3: cross-correlation statistics

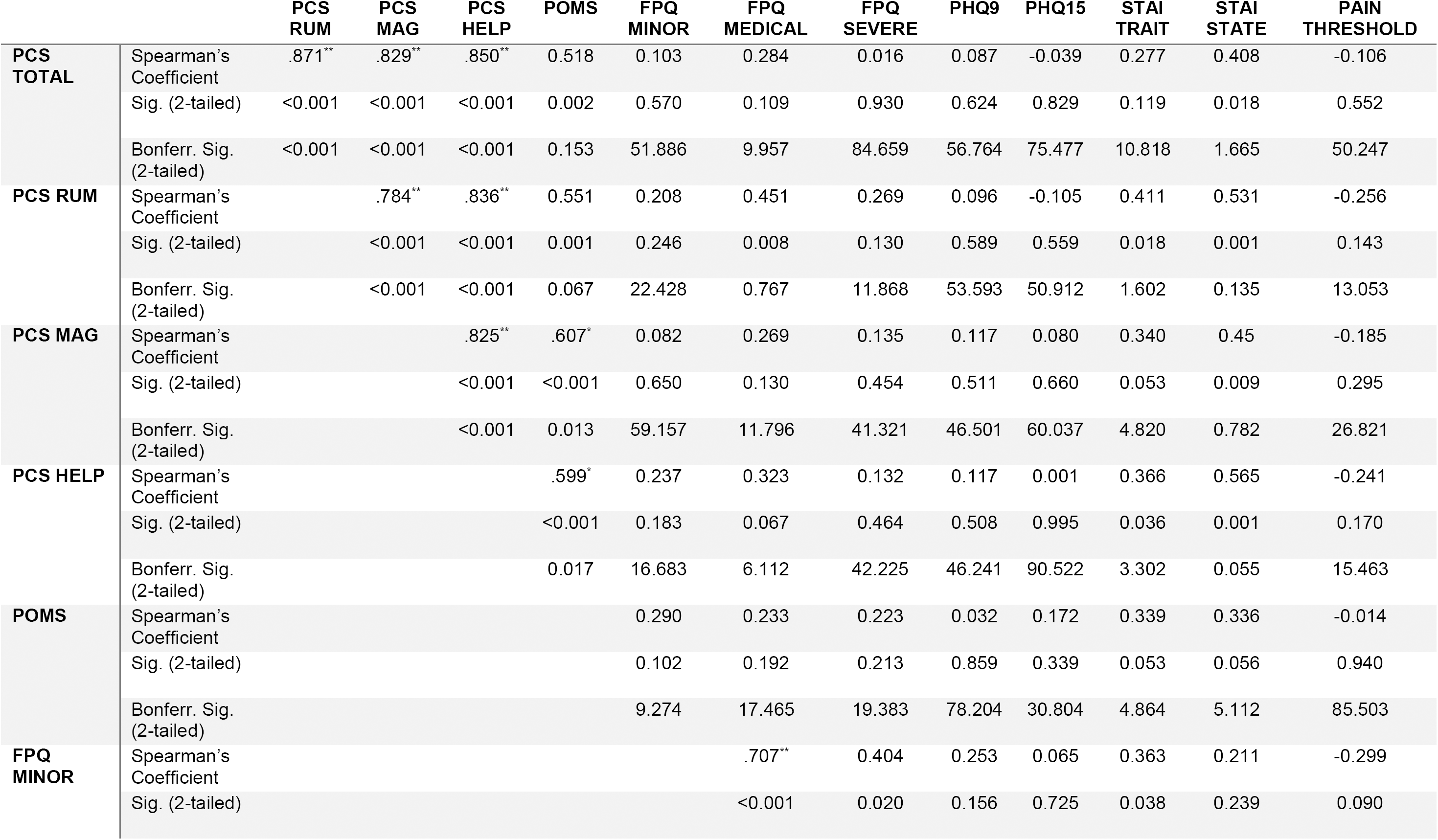
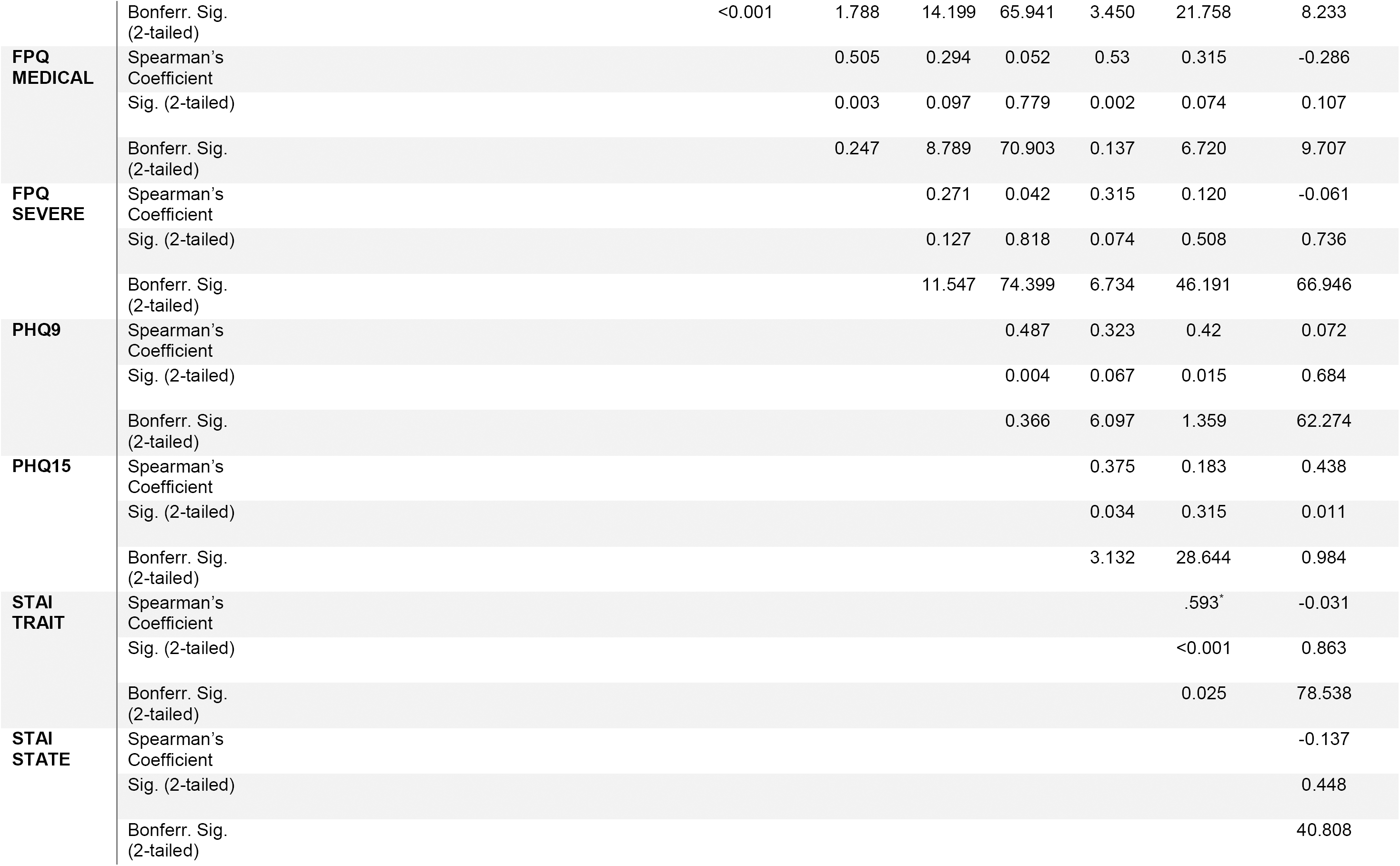
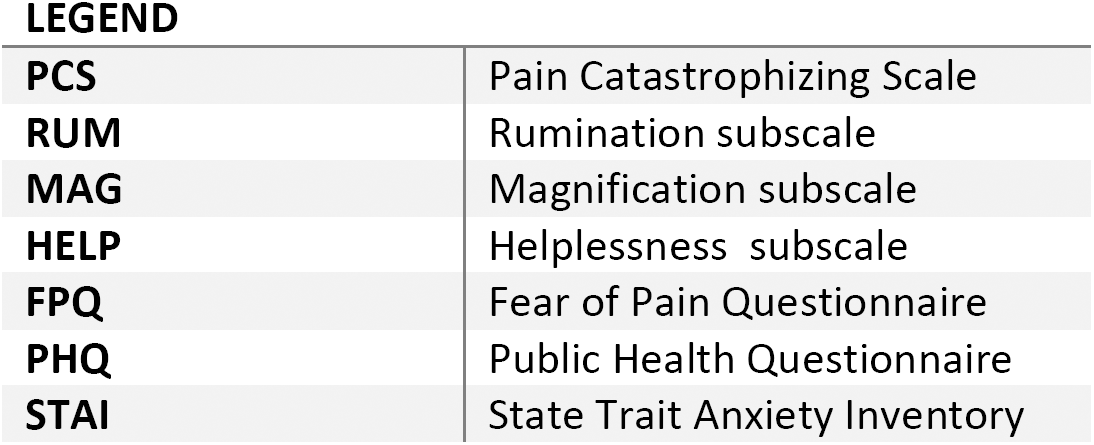

### 9.4 Supplementary Table 4: Mean pain ratings per participant/condition across trials

| SUBJECT ID | GROUP | NUMERICAL RATINGS OF PAIN /10 |  |  |  |  |  |  |  |  |
| --- | --- | --- | --- | --- | --- | --- | --- | --- | --- | --- |
|  |  | Block 1 |  |  |  | Block 2 |  |  |  |  |
|  |  | Low laser intensity |  | Medium laser intensity |  | Low laser intensity |  | Medium laser intensity |  | High laser intensity |
|  |  | Low cue | Prior high cue | Low cue | Prior high cue | Low cue | Prior high cue | Low cue | Prior high cue | High cue |
| 1 | High PC | 1.70 | 1.78 | 2.95 | 3.75 | 1.89 | 2.15 | 3.50 | 3.10 | 5.63 |
| 2 | High PC | 2.32 | 2.59 | 4.10 | 4.65 | 2.38 | 2.62 | 3.55 | 4.15 | 5.66 |
| 3 | High PC | 2.05 | 2.03 | 3.85 | 3.85 | 3.16 | 3.26 | 3.90 | 3.90 | 5.78 |
| 4 | High PC | 2.39 | 2.76 | 4.25 | 4.45 | 2.65 | 2.87 | 4.30 | 4.20 | 5.22 |
| 5 | High PC | 1.86 | 1.41 | 3.80 | 4.45 | 2.16 | 2.23 | 3.40 | 3.35 | 6.24 |
| 6 | High PC | 0.95 | 0.95 | 1.60 | 1.65 | 1.11 | 1.15 | 1.80 | 1.90 | 4.37 |
| 7 | High PC | 2.32 | 2.65 | 4.65 | 5.50 | 2.22 | 2.51 | 4.20 | 4.50 | 6.56 |
| 8 | High PC | 2.30 | 2.38 | 3.10 | 3.15 | 2.30 | 2.44 | 3.30 | 3.65 | 5.54 |
| 9 | High PC | 2.95 | 3.03 | 4.35 | 5.00 | 3.30 | 3.33 | 4.20 | 4.55 | 6.46 |
| 10 | High PC | 3.30 | 2.92 | 2.75 | 3.25 | 3.76 | 3.72 | 2.70 | 2.90 | 3.68 |
| 11 | High PC | 2.20 | 2.73 | 4.65 | 4.45 | 2.62 | 2.87 | 3.95 | 3.65 | 5.93 |
| 12 | High PC | 2.80 | 3.22 | 4.50 | 4.40 | 2.92 | 2.87 | 4.75 | 4.60 | 6.49 |
| 13 | High PC | 2.34 | 2.45 | 3.81 | 4.06 | 2.40 | 2.52 | 3.61 | 3.76 | 5.65 |
| 14 | High PC | 2.23 | 2.11 | 3.25 | 3.65 | 1.78 | 2.00 | 4.70 | 4.20 | 5.46 |
| 15 | High PC | 2.77 | 3.54 | 4.75 | 4.75 | 0.41 | 0.62 | 2.15 | 3.05 | 5.56 |
| 16 | High PC | 3.02 | 2.59 | 4.85 | 5.00 | 3.00 | 3.00 | 4.45 | 4.45 | 6.17 |
| MEAN (GROUP) |  | 2.34 | 2.45 | 3.83 | 4.13 | 2.38 | 2.51 | 3.65 | 3.74 | 5.65 |
| SD (GROUP) |  | 0.57 | 0.67 | 0.89 | 0.92 | 0.83 | 0.78 | 0.85 | 0.74 | 0.76 |
| 17 | Low PC | 1.98 | 1.84 | 4.05 | 4.60 | 2.74 | 2.87 | 4.25 | 3.94 | 6.16 |
| 18 | Low PC | 1.71 | 1.65 | 4.11 | 4.72 | 1.43 | 1.44 | 4.55 | 4.50 | 5.95 |
| 19 | Low PC | 2.98 | 2.92 | 5.20 | 5.65 | 3.00 | 2.90 | 4.25 | 4.85 | 5.66 |
| 20 | Low PC | 3.02 | 3.32 | 5.15 | 5.35 | 2.65 | 2.69 | 4.65 | 4.55 | 6.17 |
| 21 | Low PC | 1.93 | 2.49 | 3.20 | 3.60 | 2.30 | 2.49 | 2.75 | 3.35 | 5.05 |
| 22 | Low PC | 1.82 | 2.16 | 4.25 | 3.85 | 2.14 | 2.33 | 3.50 | 4.05 | 5.59 |
| 23 | Low PC | 2.23 | 1.73 | 3.70 | 3.55 | 2.43 | 2.38 | 3.35 | 3.65 | 5.93 |
| 24 | Low PC | 1.89 | 2.24 | 3.55 | 3.60 | 2.38 | 2.26 | 2.90 | 3.00 | 5.76 |
| 25 | Low PC | 3.39 | 3.49 | 4.65 | 4.70 | 3.35 | 3.56 | 4.45 | 4.35 | 6.20 |
| 26 | Low PC | 3.59 | 3.97 | 4.25 | 4.25 | 3.00 | 2.97 | 3.10 | 3.20 | 5.41 |
| 27 | Low PC | 2.91 | 3.11 | 4.65 | 3.95 | 2.27 | 2.46 | 3.35 | 4.05 | 6.22 |
| 28 | Low PC | 2.36 | 2.19 | 2.70 | 3.75 | 2.95 | 3.21 | 3.10 | 3.80 | 5.44 |
| 29 | Low PC | 1.15 | 1.05 | 1.90 | 2.15 | 1.05 | 1.38 | 2.15 | 2.10 | 3.33 |
| 30 | Low PC | 2.02 | 2.19 | 3.70 | 3.65 | 1.62 | 2.03 | 3.35 | 3.15 | 6.44 |
| 31 | Low PC | 2.14 | 2.43 | 2.85 | 3.65 | 3.16 | 2.85 | 3.40 | 3.10 | 5.34 |
| 32 | Low PC | 1.98 | 1.97 | 3.45 | 3.90 | 2.22 | 2.28 | 3.55 | 3.80 | 6.63 |
| 33 | Low PC | 3.32 | 3.38 | 3.95 | 4.20 | 2.43 | 3.21 | 4.80 | 4.95 | 5.49 |
| 34 | Low PC | 1.77 | 2.11 | 2.90 | 3.00 | 2.32 | 2.21 | 2.90 | 3.45 | 4.98 |
| MEAN (GROUP) |  | 2.34 | 2.46 | 3.79 | 4.01 | 2.41 | 2.53 | 3.58 | 3.77 | 5.65 |
| SD (GROUP) |  | 0.69 | 0.76 | 0.87 | 0.82 | 0.60 | 0.58 | 0.75 | 0.73 | 0.74 |
| SD (POOLED) |  | 0.62 | 0.71 | 0.87 | 0.86 | 0.71 | 0.67 | 0.79 | 0.72 | 0.74 |
| INDEPENDENT SAMPLES T TEST (HIGH PC > LOW PC) |  |  |  |  |  |  |  |  |  |  |
| T STATISTIC |  | 0.00 | -0.05 | 0.12 | 0.40 | -0.14 | -0.08 | 0.29 | -0.10 | -0.01 |
| P VALUE |  | 0.999 | 0.963 | 0.906 | 0.692 | 0.889 | 0.936 | 0.776 | 0.923 | 0.991 |
| EFFECT SIZE (COHEN'S D) |  | 0.00 | 0.02 | 0.04 | 0.14 | 0.05 | 0.03 | 0.10 | 0.03 | 0.00 |

### 9.5 Supplementary Table 3: mixed anova for pain ratings

| SOURCE | F | P | PARTIAL ETA SQUARED |
| --- | --- | --- | --- |
| BLOCK | 1.10 | 0.302 | 0.03 |
| BLOCK * GROUP | 0.04 | 0.846 | 0.00 |
| INTENSITY | 161.70 | <0.001 | 0.83 |
| INTENSITY * GROUP | 0.10 | 0.754 | 0.00 |
| CUE | 37.67 | <0.001 | 0.54 |
| CUE * GROUP | 0.01 | 0.938 | 0.00 |
| BLOCK * INTENSITY | 8.69 | 0.006 | 0.21 |
| BLOCK * INTENSITY * GROUP | 0.02 | 0.890 | 0.00 |
| BLOCK * CUE | 1.07 | 0.308 | 0.03 |
| BLOCK * CUE * GROUP | 0.66 | 0.422 | 0.02 |
| INTENSITY * CUE | 2.40 | 0.131 | 0.07 |
| INTENSITY * CUE * GROUP | 0.01 | 0.910 | 0.00 |
| BLOCK * INTENSITY * CUE | 1.13 | 0.296 | 0.03 |
| BLOCK * INTENSITY * CUE * GROUP | 0.74 | 0.396 | 0.02 |
| GROUP | 0.01 | 0.930 | 0.00 |

### 9.6 Supplementary Table 4: EEG sensor statistics

| CONTRAST | CLUSTER CODE | TEMPORAL EXTENT (MS) |  | P(FWE-CORR) | CLUSTER SIZE (VOXELS) | SPATIAL COORDINATES (MM) |  |
| --- | --- | --- | --- | --- | --- | --- | --- |
|  |  | Min | Max |  |  | x | y |
| **<u>VISUAL CUE PROCESSING</u>** |  |  |  |  |  |  |  |
| CUE | ss1 | 116 | 152 | 0.003 | 3303 | 9 | 56 |
| CUE | ss2 | 150 | 150 | <0.001 | 4629 | -17 | -62 |
| CUE | ss3 | 362 | 1312 | <0.001 | 127902 | 4 | -62 |
| CUE | ss4 | 384 | 1112 | <0.001 | 94700 | -68 | 18 |
| GROUP | ss5 | 256 | 604 | <0.001 | 28786 | -21 | -41 |
| GROUP | ss6 | 242 | 502 | <0.001 | 8616 | -68 | 18 |
| GROUP*CUE | ss7 | 1334 | 1440 | 0.001 | 3936 | 30 | -36 |
| **<u>NOCICEPTIVE PROCESSING</u>** |  |  |  |  |  |  |  |
| INTENSITY | ss8 | 308 | 310 | <0.001 | 8572 | -17 | 2 |
| INTENSITY | ss9 | 388 | 726 | <0.001 | 55016 | -4 | -25 |
| INTENSITY | ss10 | 382 | 670 | <0.001 | 16672 | -68 | -41 |
| EXPECTATION | ss11 | 416 | 478 | 0.002 | 2802 | 21 | -3 |

FWE: Family-wise error rate

### 9.7 Supplementary Table 5: EEG source statistics

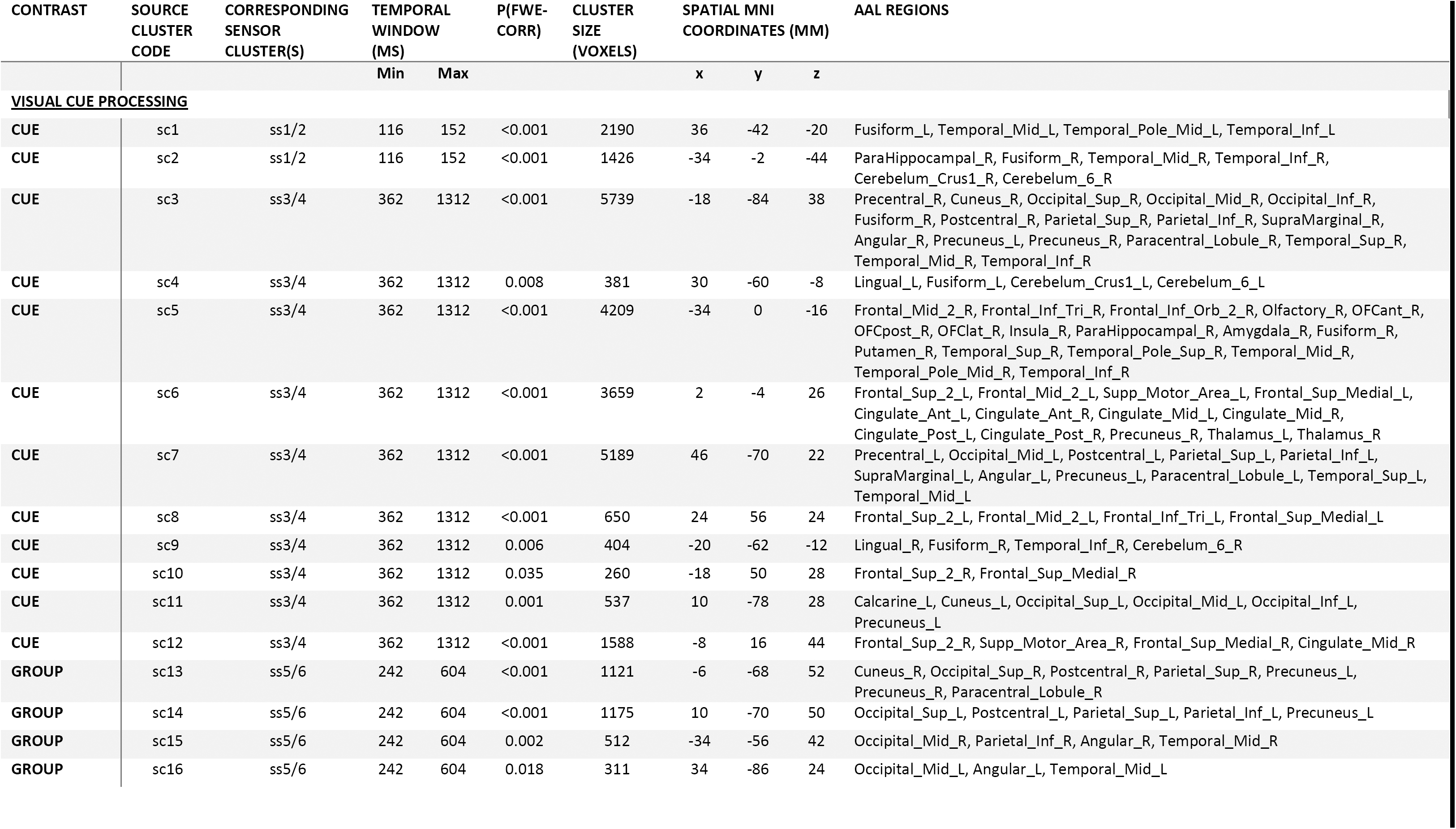
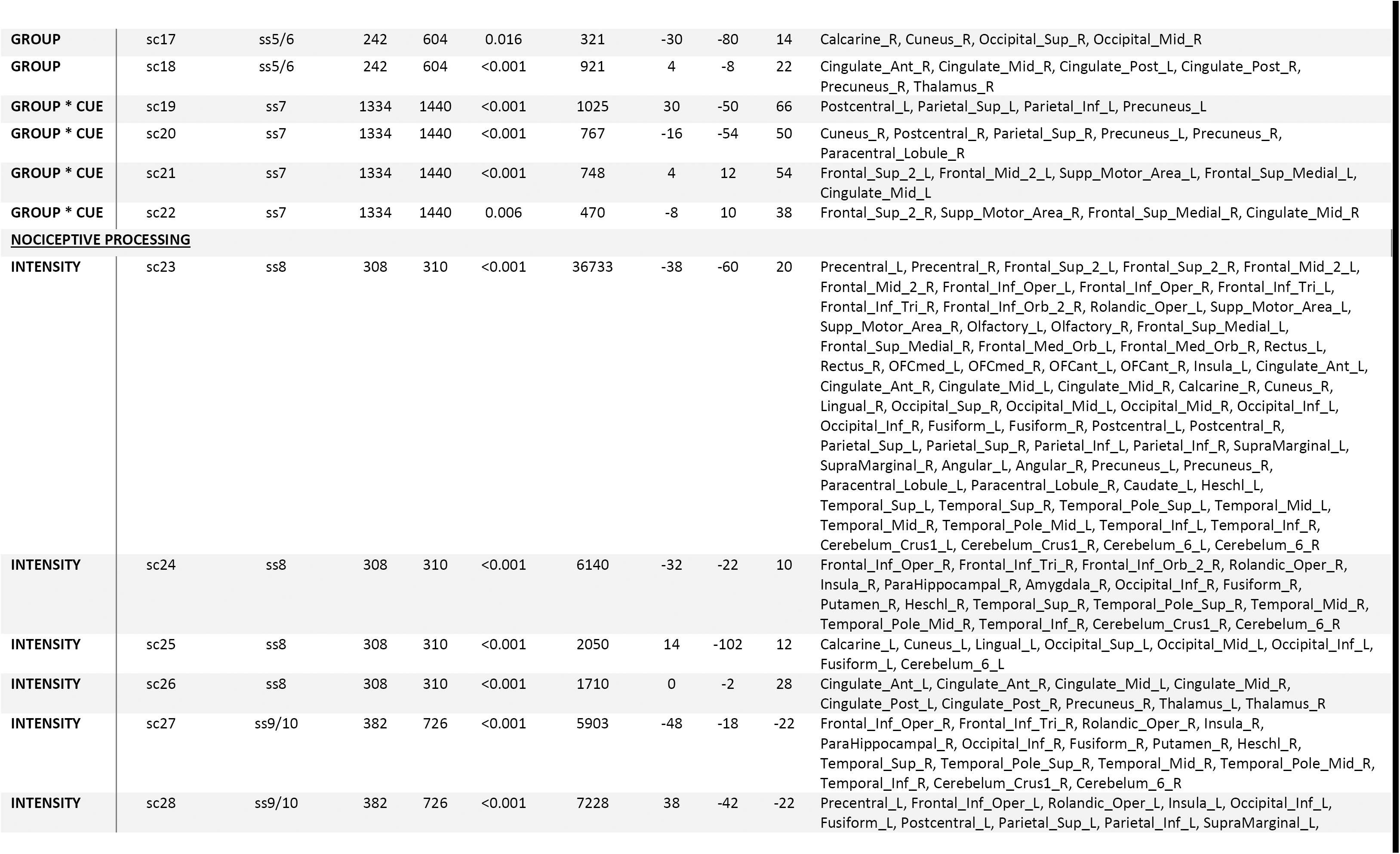
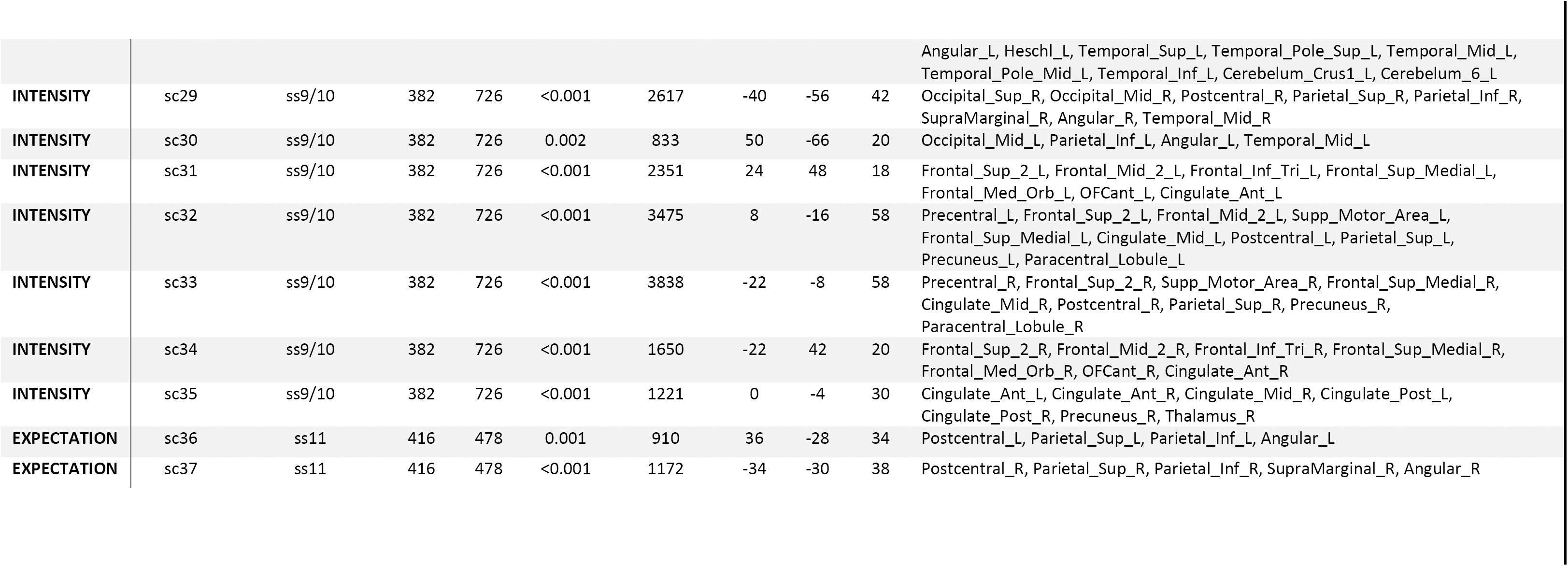

